# Tafazzin Mediates Tamoxifen Resistance by Regulating Cellular Phospholipid Composition in ER-Positive Breast Cancer

**DOI:** 10.1101/2023.08.15.553336

**Authors:** Xuan Li, Yuan Zhang, Tengjiang Zhang, Luyang Zhao, Christopher G. Lin, Haitian Hu, Hanqiu Zheng

**Affiliations:** Department of Basic Medical Sciences, School of Medicine, Tsinghua University, Beijing, 100084, China; Tsinghua-Peking Center for Life Sciences, Beijing, China; State Key Laboratory of Molecular Oncology and Department of Basic Medical Sciences, School of Medicine, Tsinghua University, Beijing, 100084, China

**Author notes:** Corresponding author **Correspondence**: Hanqiu Zheng.

**Keywords:** breast cancer, estrogen receptor-positive, tamoxifen resistance, TAZ, cardiolipin

## Abstract

Tamoxifen is the frontline therapeutic agent for the estrogen receptor-positive (ER+) subtype of breast cancer patients, which accounts for 70-80% of total breast cancer incidents. However, clinical resistance to tamoxifen has become increasingly common, highlighting the need to identify the underlying cellular mechanisms. In our study, we employed a genome-scale CRISPR-Cas9 loss-of-function screen and validation experiments to discover that Tafazzin (TAZ), a mitochondrial transacylase, is crucial for maintaining the cellular sensitivity of ER+ breast cancer cells to tamoxifen and other chemotherapies. Mechanistically, we found that cardiolipin, whose synthesis and maturation rely on TAZ, is required to maintain cellular resistance to tamoxifen. Loss of metabolic enzymatic activity of TAZ causes ERα downregulation and therapy resistance. Interestingly, we observed that TAZ deficiency also led to the upregulation of lysophosphatidylcholine (LPC), which in turn suppressed ERα expression and nuclear localization, thereby contributing to tamoxifen resistance. LPC is further metabolized to lysophosphatidic acid (LPA), a bioactive molecule that supports cell survival. Thus, our findings suggest that the depletion of TAZ promotes tamoxifen resistance through an LPC-LPA phospholipid synthesis axis, and targeting this lipid metabolic pathway could restore cell susceptibility to tamoxifen treatment.

## Introduction

Breast cancer is one of the most threatening diseases among women in recent years [1, 2]. Among breast cancer patients, about 70-80% are the estrogen receptor-positive (ER+) subtype [3, 4]. Endocrine therapy is commonly used for this subtype, targeting ERα signaling to reduce cancer cell viability. Tamoxifen, a selective ER modulator, is the primary and the frontline endocrine therapeutic agent for ER+ breast cancer [5-8]. Tamoxifen has been shown to significantly decrease the recurrence and mortality of breast cancer. However, approximately 30-40% of patients develop tamoxifen resistance, leading to therapy failure [9-11]. Overcoming tamoxifen resistance is a critical challenge in clinical practice. Many mechanisms leading to tamoxifen resistance have been reported, including alterations of ERα, crosstalk between ERα signaling and other pathways, cell cycle regulation, and protective autophagy [12]. Despite intense research, the molecular mechanism leading to tamoxifen resistance is still not fully understood, and overcoming this resistance remains a significant clinical challenge [13].

Systematic searches for genes responsible for tamoxifen resistance have been undertaken using RNA interference screens, which identify a set of novel determinants of tamoxifen sensitivity including components of a neddylation complex, modulators of chromatin remodeling, members of RAS signaling cascade, and insulin-like growth factor binding protein [14, 15]. In recent years, the development of gene editing technology, especially the CRISPR/Cas9 (Clustered regularly interspaced short palindromic repeats/Cas9) system, provides convenient tools to conduct genome-wide knockout genetic screens to identify drug resistance genes [16-22]. Notably, an epigenome screen via CRISPR/Cas9 knockout system has successfully uncovered ARID1A, a subunit of SWI/SNF chromatin remodeling complex, mediating cell sensitivity to the ER degrader fulvestrant, another widely used endocrine therapy [21]. Here, we performed a loss-of-function CRISPR/Cas9 screen using a genome-wide sgRNA library to identify genes associated with tamoxifen resistance.

Through the screen, we discovered that the expression of Tafazzin (TAZ, Gene ID: 6901. Please notice that this is not the TAZ gene involved in Hippo pathway, the gene ID of which is 25937) is a necessity to maintain cellular sensitivity to tamoxifen treatment. *TAZ* is an X-linked gene encoding a mitochondrial transacylase required for the production of cardiolipin (CL), a specific phospholipid found in mitochondria [23-27]. The metabolism of CL involves two steps: synthesis and remodeling, which are catalyzed by enzymes in the cytoplasm and inner mitochondrial membrane. TAZ is responsible for remodeling CL, upon which the vast majority of mature cardiolipin molecules depend. However, other enzymes, including ALCAT1 (acyl-CoA:lysocardiolipin acyltransferase 1) and MLCAT1 (monoacylglycerol acyltransferase 1), are also involved in CL maturation. TAZ deficiency leads to the accumulation of its substrate monolysocardiolipin (MLCL), and the reduction of the CL [25, 28]. Our results reveal that several phospholipids are also essential in generating tamoxifen resistance.

## Results

### Genome-wide CRISPR-Cas9 screen for identifying tamoxifen resistance genes

We applied a genome-wide CRISPR-Cas9 knockout screening to identify genes involved in therapeutic resistance to tamoxifen therapy, using the commonly used ER+ breast cancer cell line MCF7 as the model system. MCF7 cells were transduced with the Brunello sgRNA library in lentiCRISPR v2 (one vector system) of 76,441 small guide RNAs (sgRNAs), which targets to knockout 19,114 genes. [29, 30]. One thousand non-targeting sgRNAs were included in this library. Tumor cells were selected by tamoxifen citrate (shortened as tamoxifen in the following text) treatment for three days and the surviving cells were considered to be drug-resistant. Genomic DNA was isolated from these cells, and next-generation sequencing analysis was performed (Fig. 1A). The abundance of each sgRNA in the original library was determined, with most sgRNAs being evenly distributed and few being either low or high in abundance (Supplementary Fig. S1A). We analyzed the enrichment or depletion of sgRNAs by comparing the tamoxifen-treated group to the DMSO-treated group, and the differential sgRNA presentation in each group was determined using a Model-based Analysis of the Genome-wide CRISPR/Cas9 Knockout (MAGeCK) algorithm [31]. We obtained nine candidate genes by setting a threshold with a false discovery rate (FDR) of less than 10% and a fold change greater than 2 folds (Fig. 1B). These candidate genes were: *RPS16, PELO, USPL1, CCDC81, NDUFB4, IMP3, PSMD13, ROMO1,* and *TAZ*. Additionally, the other leading increasing or decreasing genes were showed and some of these genes might also be related to tamoxifen resistance (Supplementary Fig. S1B&C).

**Figure 1.**
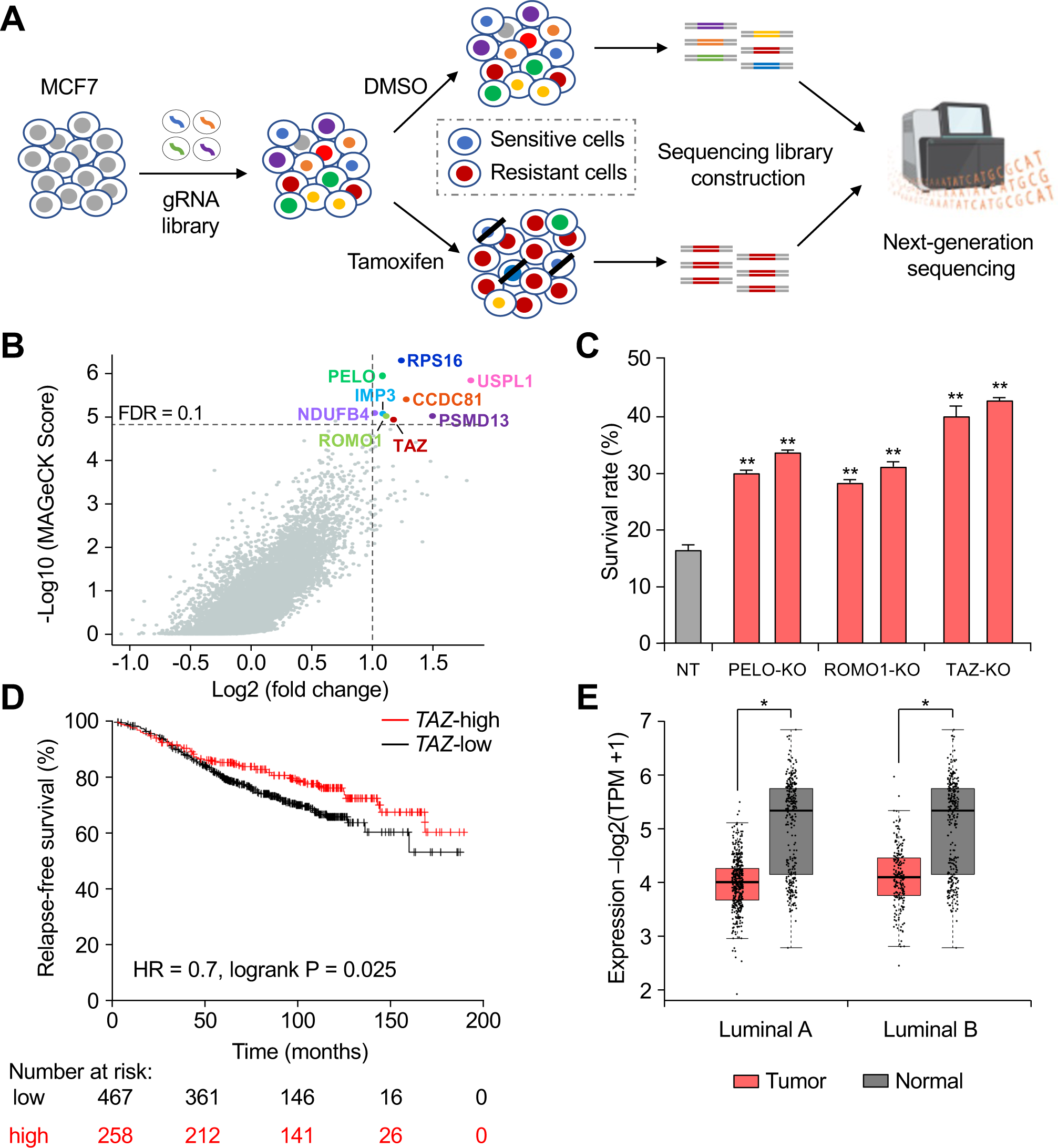
Genome-wide screen for tamoxifen resistance genes in ER-positive breast cancer cells. **A.** The screening workflow. MCF7 cells were transduced with one vector sgRNA library and puromycin selected to generate stable cells with the expected MOI of 0.2. Cells were then exposed to either DMSO or tamoxifen treatment, surviving cells were lysed for DNA isolation and DNA sequencing library construction for the next-generation sequencing. **B.** Candidate genes were plotted based on mean log2 fold change of sgRNA counts compared to control selection (DMSO) and -log10 score computed by MaGeCK. The transverse dashed line indicates an FDR (false discovery rate) of 0.1 and the vertical dashed line indicates an LFC (log2 fold change) of 1.0. Annotated genes were the top candidate genes. **C.** Candidate gene KO cell lines were constructed in MCF7 cells, including 1 control sgRNA (NT), and 2 independent sgRNAs for each candidate gene. Cells were treated with DMSO or 12 µM tamoxifen for 48 h. Data represents the cell viability normalized to the corresponding DMSO control group. n = 3, Data presented as mean ± SD. ** p < 0.01, by Student’s t-test. **D.** Kaplan-Meier survival curves suggest that the lower mRNA expression of *TAZ* is correlated with poor patient relapse-free survival (RFS) with tamoxifen based on the datasets of ER array positive patients following only tamoxifen therapy in https://kmplot.com. **E.** The mRNA expression level of *TAZ* is lower in breast tumor tissues compared to normal tissues in both Luminal A and Luminal B subtypes of breast cancers. Data was analyzed based on TCGA and GTEx data set on GEPIA2 (http://gepia2.cancer-pku.cn). * p < 0.05. TPM, Transcripts per million. **See also Supplemental Figure S1.**

To confirm the screening results, MCF7 cells were labeled with *firefly luciferase*, and subsequent experiments measured cell survival using in vitro luciferase assay. Stable knockout (KO) cell lines were generated for the target genes using the two best performing sgRNAs for each gene. The knockout of RPS16, which encodes a ribosomal protein, proved toxic to the cells. This is consistent with data from Cancer Dependency Map (DepMap) that indicates cells rely on RSP16 for survival (data not shown). For these candidate gene KO cells, the indel frequencies were determined by the method of Tracking Indels by Decomposition (TIDE) to confirm gene knockout efficiencies (Supplementary Table S5) [32].

To validate the phenotype of the above KO cells, we assessed the survival rates of each group following tamoxifen treatment compared to Non-Target (NT) control cells. Using two independent sgRNAs, we observed significant increases in cell survival rates after tamoxifen treatment for the PELO, ROMO1, and TAZ KO groups, whereas the KO of other genes demonstrated no significant, or inconsistent survival benefits between two sgRNA KO cell strains (Fig. 1C and Supplementary Fig. S1D). To further investigate the clinical relevance of these three genes, we performed patient survival analysis by using an online KMplot algorism, which is based on multiple patient datasets. We found that lower expression level of *ROMO1* or *TAZ* is associated with poorer patient relapse-free survival (RFS) in patients undergoing tamoxifen therapy (Fig. 1D and Supplementary Fig. S1E). Moreover, even in patient dataset without considering their treatment history, lower expression level of *TAZ* is still correlated with poorer patient RFS of ER+ breast cancer (Supplementary Fig. S1F). In contrast, lower expression level of PELO is correlated with better patient relapse-free survival (Supplementary Fig. S1G). Additionally, only TAZ was significantly downregulated in ER+ breast tumor tissues compared to normal tissues, while ROMO1 had a tendency of higher expression in tumor tissues than in normal tissues, although it did not reach statistical significance (Fig. 1E and Supplementary Fig. S1H). These findings suggest that TAZ may play a critical role in maintaining the sensitivity to tamoxifen treatment, and its loss of expression may contribute to the development of tamoxifen resistance in ER+ breast cancer patients.

Although there have been numerous studies exploring the mechanisms of resistance to endocrine therapy for breast cancer [10, 33-37], the potential impact of intracellular phospholipid metabolism on tamoxifen resistance remains largely unknown. Our findings show that TAZ KO results in therapeutic resistance to tamoxifen, coupled with the fact that TAZ encodes an enzyme involved in mitochondrial cardiolipin metabolism [23, 38], suggest that investigating the functional role of TAZ in tamoxifen resistance and exploring the potential impact of intracellular phospholipid metabolism on tamoxifen resistance could be valuable research directions.

### Depletion of TAZ induces tamoxifen resistance

To confirm that TAZ depletion induces tamoxifen resistance, we generated TAZ knockout cell lines using three independent sgRNAs in two additional ER+ breast cancer cell lines, T47D and ZR-75-1. We confirmed the KO efficiencies of TAZ using TIDE analysis (Supplementary Figs. S2A-C). KO of TAZ increased cell viability compared to NT cells at two different concentrations of tamoxifen treatment in both cell lines (Figs. 2A&B). In MCF7 cells, TAZ depletion increased the number of colonies during tamoxifen treatment in a long-term colony formation assay and led to an increased number of tumor spheres in a tumor sphere formation assay (Figs. 2C&D and Supplementary Fig. S2D). We then tested our findings *in vivo* by following experimental procedures shown in Fig. 2E. Briefly, we implanted 4-6 weeks old female nude mice subcutaneously with a slow-releasing estrogen pellet, followed by mammary fat pad injection with 2 × 10^6^ vector control or TAZ-KO MCF7 cells. Tumor volume was measured every four days. Mice were then treated with tamoxifen three times a week after the tumor volume reached 60 mm^3^, and continued until the experimental endpoint (Fig. 2E). Tamoxifen administration significantly reduced the tumor volume of control (NT) xenografts, whereas TAZ-deficient cells were extremely resistant to tamoxifen and showed no obvious growth inhibition (Fig. 2F and Supplementary Fig. S2E).

**Figure 2.**
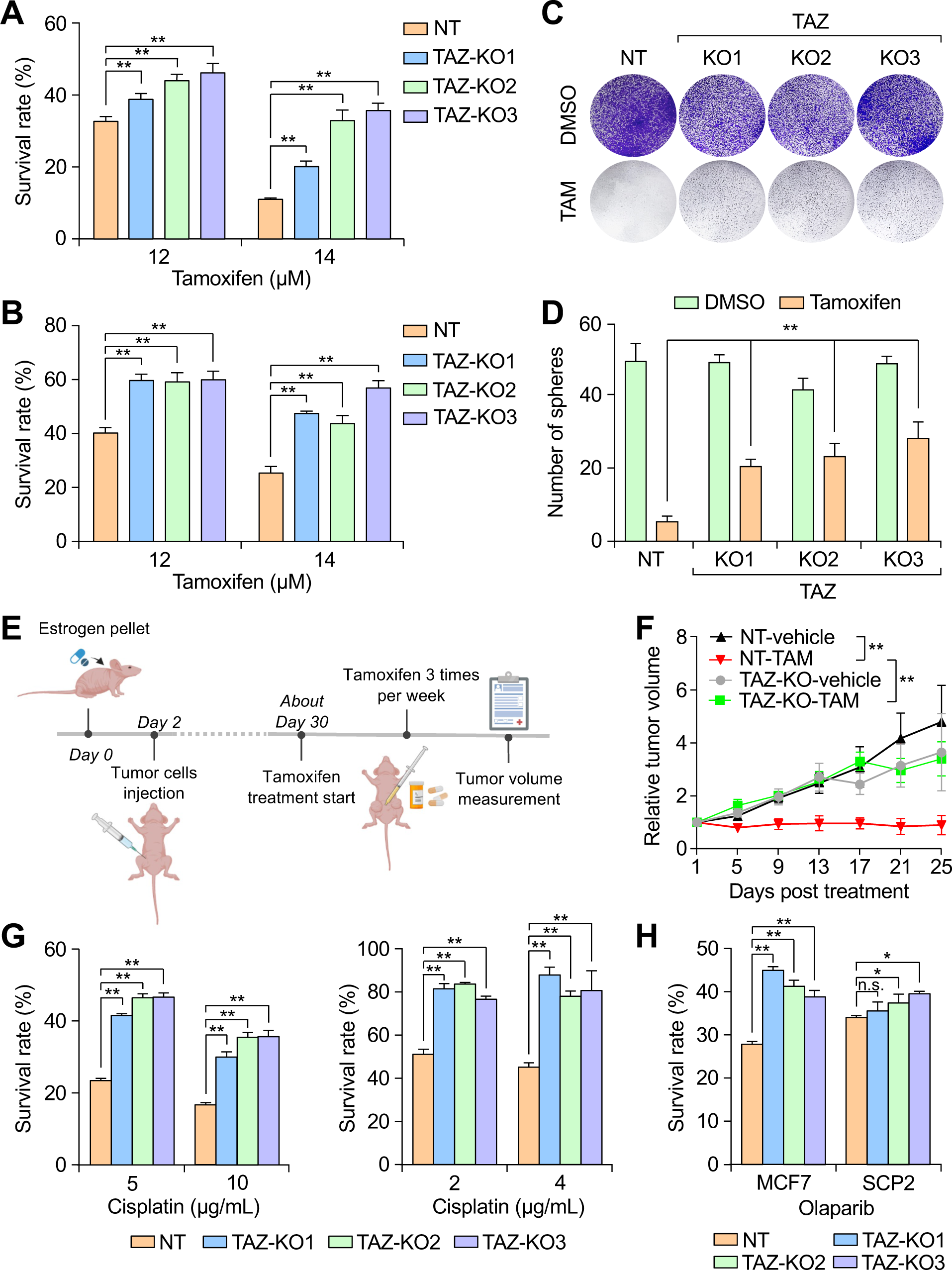
Depletion-of-TAZ induces multiple drug resistance in ER-positive breast cancer. **A, B.** TAZ-KO in T47D (A), or in ZR-75-1 (B) cells, or NT control cells were treated with DMSO or indicated concentrations of tamoxifen for 48 h. Data represents the cell viability normalized to the corresponding DMSO control group. n = 3 per group, data presented as mean ± SD. ** p < 0.01, by Student’s t-test. **C.** The colonies of NT or TAZ-KO MCF7 cells were treated with DMSO or 14 µM tamoxifen and stained by crystal violet. **D.** Tumor spheres of NT or TAZ-KO MCF7 cells were treated with DMSO or 10 µM tamoxifen and the number of spheres was counted. n = 3 per group, data presented as mean ± SD. ** p < 0.01, by Student’s t-test. **E.** The schematic illustration of MCF7 xenograft experimental procedure. 2 × 10^6^ TAZ-KO MCF7 or NT control cells were injected into the mammary fat pads of 6 weeks old female nude mice. When tumor volume reached around 60 mm^3^, mice were treated with either vehicle control or tamoxifen three times a week until experimental endpoint. The tumor volume was measured every four days. **F.** Quantification of tumor volume from experiment in E. TAZ-KO-DMSO group: n = 6, TAZ-KO-TAM group n= 8, NT-DMSO group n = 5, NT-TAM group n = 6. Data presented as mean ± SEM. ** p < 0.01 by two-way ANOVA. **G.** TAZ-KO MCF7 (left) and ZR75-1 (right), or their respective NT control cells were treated with either PBS or indicated concentrations of cisplatin for 24 h. Data represents the cell viability normalized to the corresponding PBS control group. n = 3 per group, data presented as mean ± SD. ** p < 0.01 by Student’s t-test. **H.** TAZ-KO MCF7 and SCP2 cells, as well as respective NT control cells were treated with DMSO or 50 μM olaparib for 72 h. Data represents the cell viability normalized to the corresponding DMSO group. n = 3 per group, data presented as mean ± SD. * p < 0.05, ** p < 0.01, n.s. represents no significant difference by Student’s t-test. **See also Supplemental Figure S2.**

To determine whether TAZ deficiency causes resistance to a broad range of therapeutic agents, we treated MCF7 and ZR-75-1 with cisplatin, a chemotherapeutic agent that induces DNA damage, and assessed their cell survival rates. The KO of TAZ protected both cells from death caused by cisplatin (Fig. 2G). Interestingly, KO of TAZ also enhanced cell survival after treatment with Olaparib, a Poly (ADP-ribose) polymerase inhibitor (PARPi), which is typically used in DNA repair-deficient cancers (Fig. 2H). However, the KO of TAZ did not confer any obvious survival benefit during cisplatin and Olaparib treatments in MDA-MB-231 or SCP2 cells, both of which are triple-negative breast cancer cells (Fig. 2H and Supplementary Figs. S2F-I). These findings suggest that TAZ depletion-induced drug resistance might be specific to ER+ breast cancer cells, but not to triple-negative breast cancers.

### TAZ deficiency leads to ERα downregulation and cell cycle arrest

To investigate the potential molecular mechanism underlying TAZ-mediated tamoxifen resistance, we examined the expression of the estrogen receptor α (ERα) coding gene, *ESR1*. KO of TAZ dramatically reduced *ESR1* mRNA expression (Fig. 3A), and almost completely eliminated the protein expression of ERα (Fig. 3B). A similar decrease in ERα protein expression could be seen in ZR-75-1 cells after TAZ KO (Supplementary Fig. S3A). As a transcription factor, ERα must translocate from cytoplasm to the nucleus to function properly. Therefore, we measured its intracellular localization and found that while ERα was predominantly localized to the nucleus in control cells, KO of TAZ resulted in a significant decrease in nuclear localization and a clear increase in cytoplasmic localization (Fig. 3C). Consistent with the decreased protein expression and nuclear localization of ERα, the expression levels of many classical ERα signaling genes, including *CCND1*, *GREB*, *MYC*, *PGR*, *TFF1-3*, were significantly down-regulated in TAZ KO cells both in RNA and protein levels (Fig. 3D, and Supplementary Fig. S3B). Cell cycle analysis by propidium iodide (PI) staining and fluorescence-activated cell sorting (FACS) revealed a significant decreased in the S phase and G2/M phase, and an increased G0/G1 cells in TAZ-KO cells compared to control cells (Fig. 3E). In summary, these results indicate that TAZ KO led to decreased ERα expression and nuclear localization, resulting in the reduction of downstream gene expression and slowed cell cycle progression, with a clear G1 arrest. Notably, the G1-arrest was not apparent in TAZ-KO SCP2 triple-negative breast cancer cell line (Supplementary Fig. S3C), indicating that the aforementioned drug resistances induced by TAZ KO in the ER+ cell lines may be associated with their ability to induce cell cycle arrest.

**Figure 3.**
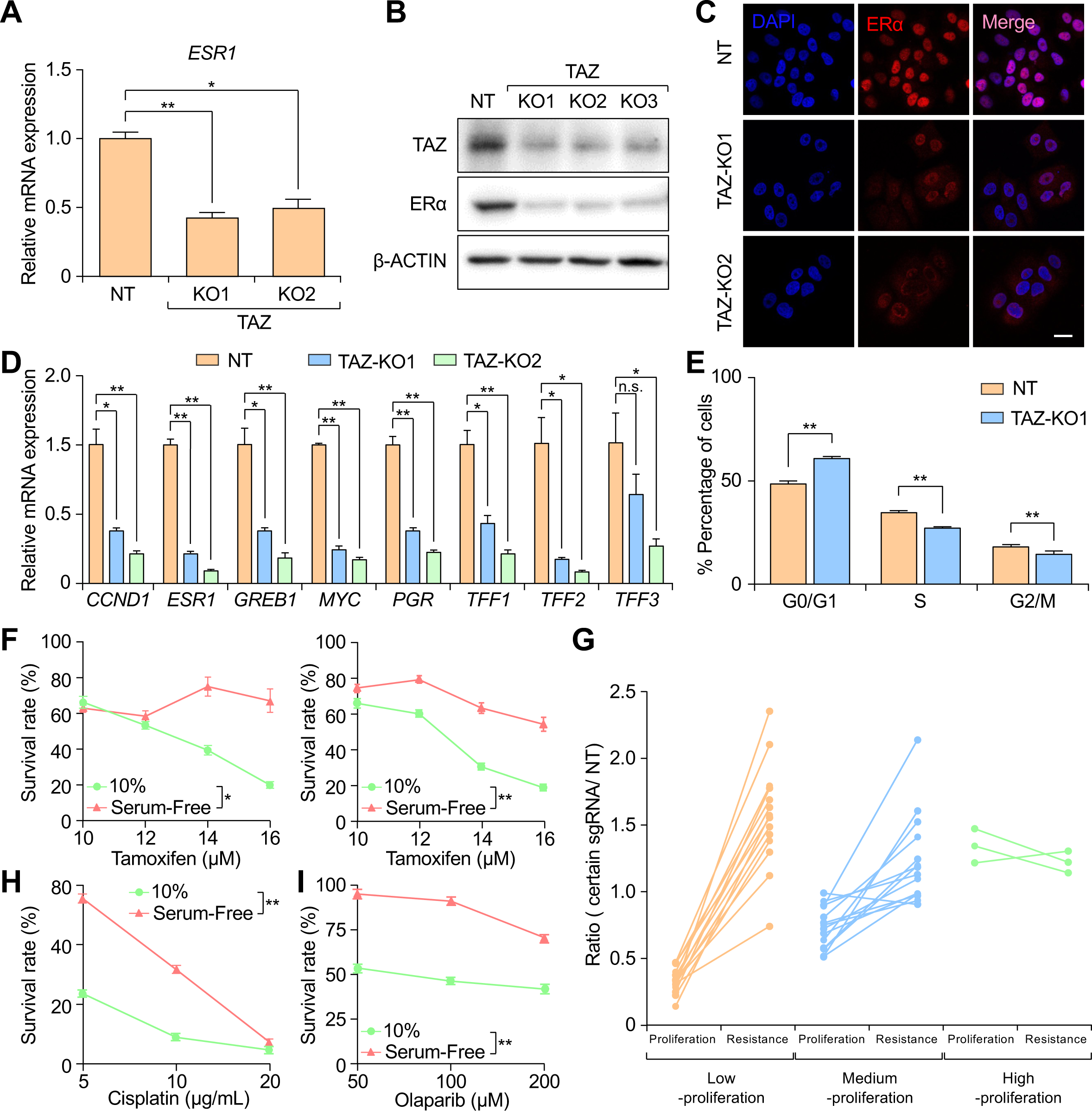
Loss-of-TAZ down-regulates ERα signaling and reduces cell proliferation rate. **A.** mRNA was isolated from NT control or TAZ-KO MCF7 cell lines. The relative *ESR1* mRNA expression levels in these cells were determined by quantitative PCR (qPCR). *GAPDH* was used as the internal control. n = 3. Data presented as mean ± SD. * p < 0.05, ** p < 0.01, by Student’s t-test. **B.** The TAZ and ERα protein levels in TAZ-KO MCF7 cells and in NT control cells were determined by immunoblotting. β-ACTIN was used as the internal loading control. **C.** NT control or TAZ-KO MCF7 cells were fixed for immunofluorescent staining against ERα (red). Nuclei were stained with DAPI (blue). Images were captured by confocal microscopy. Scale bar, 20 μm. **D.** Relative mRNA expression levels of ERα pathway downstream genes in NT control or TAZ-KO MCF7 cells were detected by qPCR. *GAPDH* was used as the internal control. n = 3, data presented as mean ± SD. n.s. represents no significant difference, * p < 0.05, ** p < 0.01, by Student’s t-test. **E.** Cell cycle analysis by PI staining and FACS analysis. n = 3, data presented as mean ± SD. ** p < 0.01, by Student’s t-test. **F.** MCF7 (left panel) or T47D (right panel) cells were cultured in medium containing 10% serum or starved in serum-free medium for 24 h, then treated with DMSO or increasing concentrations of tamoxifen for 48 h. The cell survival rate at each concentration was plotted. n = 3, data presented as mean ± SD. * p < 0.05, ** p < 0.01, by t-test. **G.** The correlation between cell proliferation index and tamoxifen resistance ratio. Each line represents a sgRNA against one specific gene. According to the proliferation index, the sgRNAs were divided into 3 groups: Low-proliferation (proliferation index ≤ 0.5), Medium-proliferation (0.5 < proliferation index < 1), and High-proliferation (proliferation index ≥ 1). **H, I.** MCF7 cells were cultured in normal medium containing 10% serum or starved in serum-free medium for 24 h, and then treated with vehicle control or increasing concentrations of cisplatin (H) or Olaparib (I) for 24 h or 72 h respectively. The cell survival rate at each concentration was plotted. n = 3, data presented as mean ± SD. * p < 0.05, ** p < 0.01, by t-test. **See also Supplemental Figure S3.**

Dormant tumor cells and recent discovered drug-tolerant persister cells with quiescent or slowly-cycling state have the capacity to escape the anti-tumor therapies and obtain long-term survival [39-41]. Slow cell cycling might also be a common trait for tamoxifen resistance cells. To investigate this hypothesis, we induced cell cycle arrest at the G1 phase in MCF7 and T47D cells by culturing them with serum-free medium, and subsequently treated them with tamoxifen (Supplementary Fig. S3D&E). The results demonstrated that cells were more resistant to tamoxifen treatment under serum-starvation conditions than under normal culture conditions (Figs. 3F). Additionally, we created stable KO strains of MCF7 using selected sgRNAs from the top 10% rank from our initial gRNA screening (Supplementary Table S2) and tested the sensitivity of tumor cells to tamoxifen treatment after transduction with individual sgRNAs, as well as their effect in slowing down cell cycle progression. A correlation between cell proliferation index and tamoxifen resistance was observed, whereby cells with lower proliferation rates (proliferation index < 0.5) were more resistant to tamoxifen treatment, while cells with mild or no suppression of cell proliferation (proliferation index > 0.5) were almost not resistant to tamoxifen treatment (Fig. 3G). Cell cycle analysis of cell lines bearing the KO of these genes further revealed that cells bearing strong tamoxifen resistance (resistance ratio > 1.5) generally had a clear cell cycle arrest at the G1 phase (Supplementary Fig. S3F). These cells with lower proliferation rates showing cell cycle arrest, which could be considered as quiescent cells, were likely to be more resistant to therapy. Moreover, serum starvation-induced cell cycle arrest also led to therapy resistance of MCF7 cells to cisplatin and Olaparib treatments (Figs. 3H&I). Interestingly, KO of TAZ resulted in a modest decrease in the cell proliferation rate but generated a significantly strong therapeutic resistance phenotype, suggesting that additional mechanisms might be involved in TAZ KO-induced resistance.

### TAZ enzymatic activity is responsible for ERα downregulation and drug resistance

To determine how TAZ affects ERα expression and induces tamoxifen resistance, we investigated whether there is a direct protein-protein interaction between TAZ and ERα. To this end, we generated HA-tagged ERα and FLAG-tagged TAZ constructs and co-transfected them into HEK293T cells for co-immunoprecipitation (co-IP) assay. However, there was no observable interaction between these two proteins (Supplementary Fig. S4A).

We investigated whether the enzymatic activity of TAZ is relevant to its role in tamoxifen resistance, given that TAZ is a mitochondrial-localized, phospholipid metabolic enzyme with transacylase activity. The H4XD motif, located in the N-terminal region of TAZ, is essential for its catalytic activity (Fig. 4A) [23]. To explore the role of this motif in tamoxifen resistance, we generated an enzyme-dead mutant lacking the H4XD motif (named TAZ-ΔH4XD) and examined its cellular localization in MCF7 cells. TAZ-ΔH4XD colocalized with MitoTracker, a mitochondrial dye, similar to the wild-type TAZ protein, suggesting that the deletion of the catalytic center did not affect TAZ’s cellular localization (Fig. 4B).

**Figure 4.**
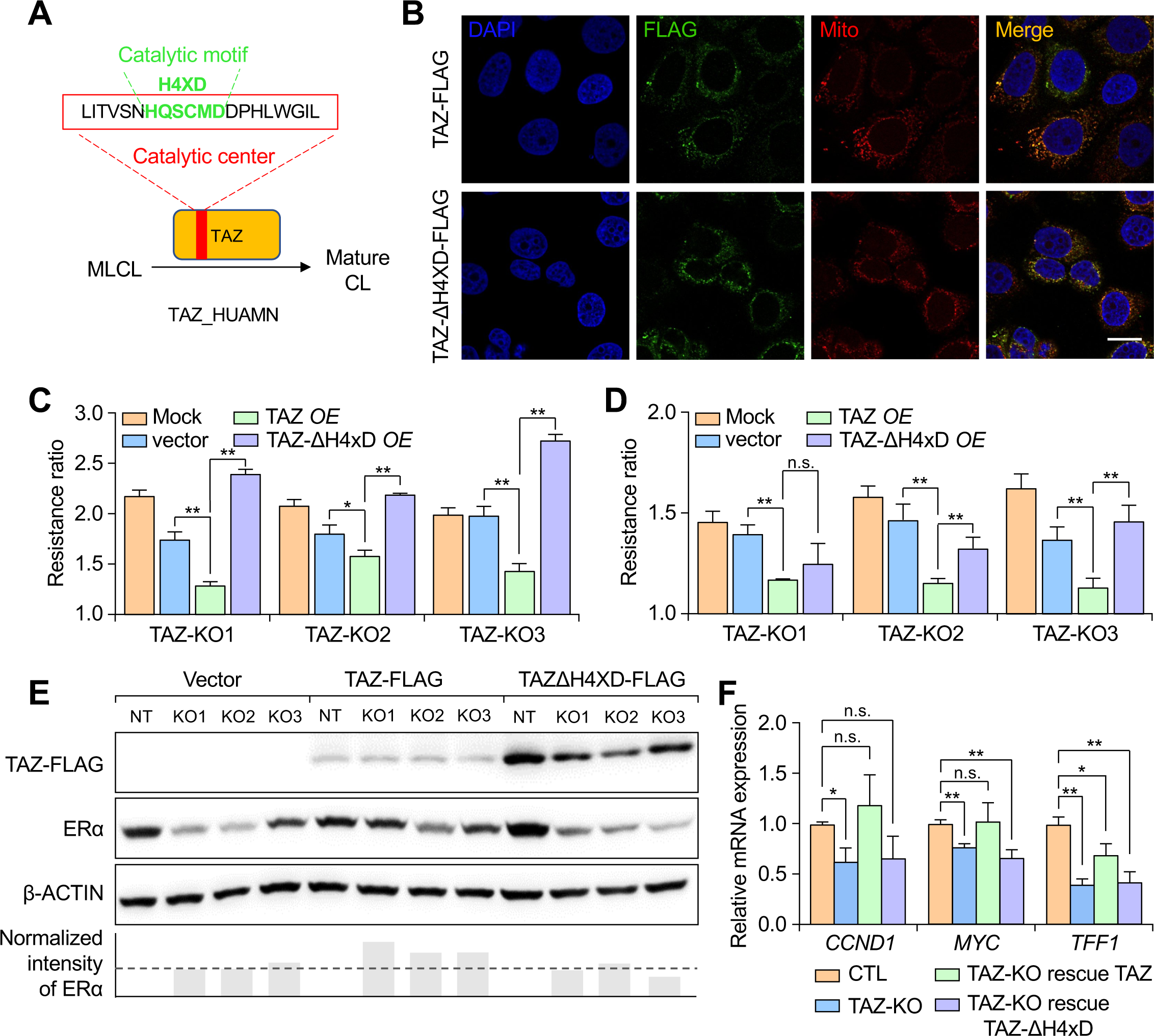
Metabolic enzymatic activity of TAZ is essential for its role in tamoxifen resistance. **A.** A Schematic illustration of catalytic center for TAZ. The red box represents the conserved catalytic canter and the green font shows the HXXXXD (H4XD) motif characteristic of acyltransferases. **B.** Stable overexpression of exogenous TAZ-FLAG or truncated TAZ-ΔH4XD-FLAG in MCF7 using lentivirus infection. The expression and localization of TAZ were detected by immunofluorescent staining using FLAG antibody (green) and Mito-tracker probe (red) respectively. Nuclei were stained with DAPI (blue). Images were captured by confocal microscopy. Scale bar, 20 μm. **C, D.** Wild type TAZ or TAZ-ΔH4XD mutant were re-expressed on TAZ-KO MCF7 cells, an empty vector was used as a control. Cells were treated with tamoxifen (C) or cisplatin (D) for 48 h. The cell survival ratio for each group was normalized to NT control group, which was not shown on the graph. n = 3, data presented as mean ± SD. * p < 0.05, ** p < 0.01, n.s. represents no significant difference by Student’s t-test. **E.** The protein expression levels of FLAG-tagged TAZ and ERα of cells from experiment in C were determined by immunoblotting. β-ACTIN was used as the internal loading control. The relative ERα expression was calculated by image densitometry analysis compared to β-ACTIN control using ImageJ software. **F.** The mRNA from NT control, TAZ-KO, TAZ-KO with wild type TAZ rescue, TAZ-KO with TAZ-ΔH4XD mutant rescue MCF7 cells were lysed for mRNA purification. The relative mRNA expression levels of several ER pathway downstream genes were determined by qPCR. *GAPDH* was used as the internal control. n = 3, data presented as mean ± SD. * p < 0.05, ** p < 0.01, n.s. represents no significant difference by Student’s t-test. **See also Supplemental Figure S4.**

We then evaluated the effect of rescue the tamoxifen resistance phenotype of TAZ-KO MCF7 cells by either wild-type TAZ or TAZ-ΔH4XD mutant. We confirmed that wild-type TAZ, but not TAZ-ΔH4XD expression could rescue cardiolipin (CL) and monolysocardiolipin (MLCL) levels in TAZ-KO cells, indicating the importance of the intact metabolic enzymatic activity of TAZ (Supplementary Fig. S4B&C). Intriguingly, TAZ overexpression significantly reduced the therapy resistance to tamoxifen caused by TAZ deficiency, while TAZ-ΔH4XD mutant still conferred resistance to tamoxifen therapy. This rescue phenotype was observed in TAZ-KO cells generated by three independent sgRNAs (Fig. 4C), and in TAZ knockdown cells generated by shRNA construct (Supplementary Fig. S4D&E). TAZ overexpression also significantly reduced the resistance to cisplatin, while TAZ-ΔH4XD mutant was unable to rescue the resistance to cisplatin treatment (Fig. 4D).

As we previously observed, TAZ KO led to a significant decrease in ERα expression. Upon re-expression of TAZ, the protein expression level of ERα was restored, while expression of TAZ-ΔH4XD could not rescue the ERα expression (Fig. 4E). TAZ re-expression, but not TAZ-ΔH4XD mutant, restored the mRNA expression of several ERα downstream genes, including *CCND1*, *MYC*, and *TFF1* (Fig. 4F). In summary, our results suggest that the enzymatic activity of TAZ is crucial for its role in tamoxifen resistance and the maintenance of ERα expression.

### A malfunction of the cardiolipin-reliant cellular mechanism induces tamoxifen resistance

Because TAZ is an essential enzyme that plays a crucial role in the maturation of cardiolipin (CL), we hypothesized that a deficiency in TAZ could result in the downregulation of ERα and drug resistance by directly impacting intracellular phospholipid metabolism. TAZ facilitates the transition of monolysocardiolipin (MLCL) to mature CL by remodeling cardiolipin (Fig. 4A). Therefore, we used high-performance liquid chromatography-mass spectrometry (HPLC-MS) to examine phospholipid metabolism levels in TAZ-knockout MCF7 cells and observed significant perturbances (Fig. 5A). As expected, we found a significant accumulation of MLCL and a marked decrease in CL levels in TAZ-KO MCF7 cells compared to NT-control cells (Fig. 5B). To confirm that the perturbances in CL metabolism were responsible for drug resistance, we further disrupted CL metabolism by knocking down the expression of other enzymes involved in CL biosynthesis, such as TAMM41, PGS1, PTPMT1, and ALCAT1 (Supplementary Figs. S5A&B). Depletion of the expression of these genes also induced cellular resistance to tamoxifen in MCF7 cells, confirming the crucial role of cardiolipin metabolism in therapy resistance (Fig. 5C). There are two types of extrinsically-induced apoptotic signaling: type I involves only the caspase-8 processing in the cytosol, whereas type II relies on the caspase-8 processing in both cytosol and mitochondria [42-44]. Previous research has shown that CL on the mitochondria membrane plays a critical role in the type II apoptotic response as a platform for caspase-8 recruitment, activation, and processing [42]. To investigate whether the abnormal CL metabolism causes tamoxifen resistance by disrupting type II apoptosis, we measured caspase-8 processing by detecting its cleavage product, p43, after tamoxifen treatment (Supplementary Fig. S5C). We found that only the caspase-8 processing on the mitochondria was blocked due to the abnormal CL metabolism induced by TAZ-KO, as evidenced by decreased p43 protein levels in TAZ-KO samples (Fig. 5D). This result suggests that the dysfunction of type II apoptosis may be partially responsible for the aforementioned downstream phenotypes of CL disorder.

**Figure 5.**
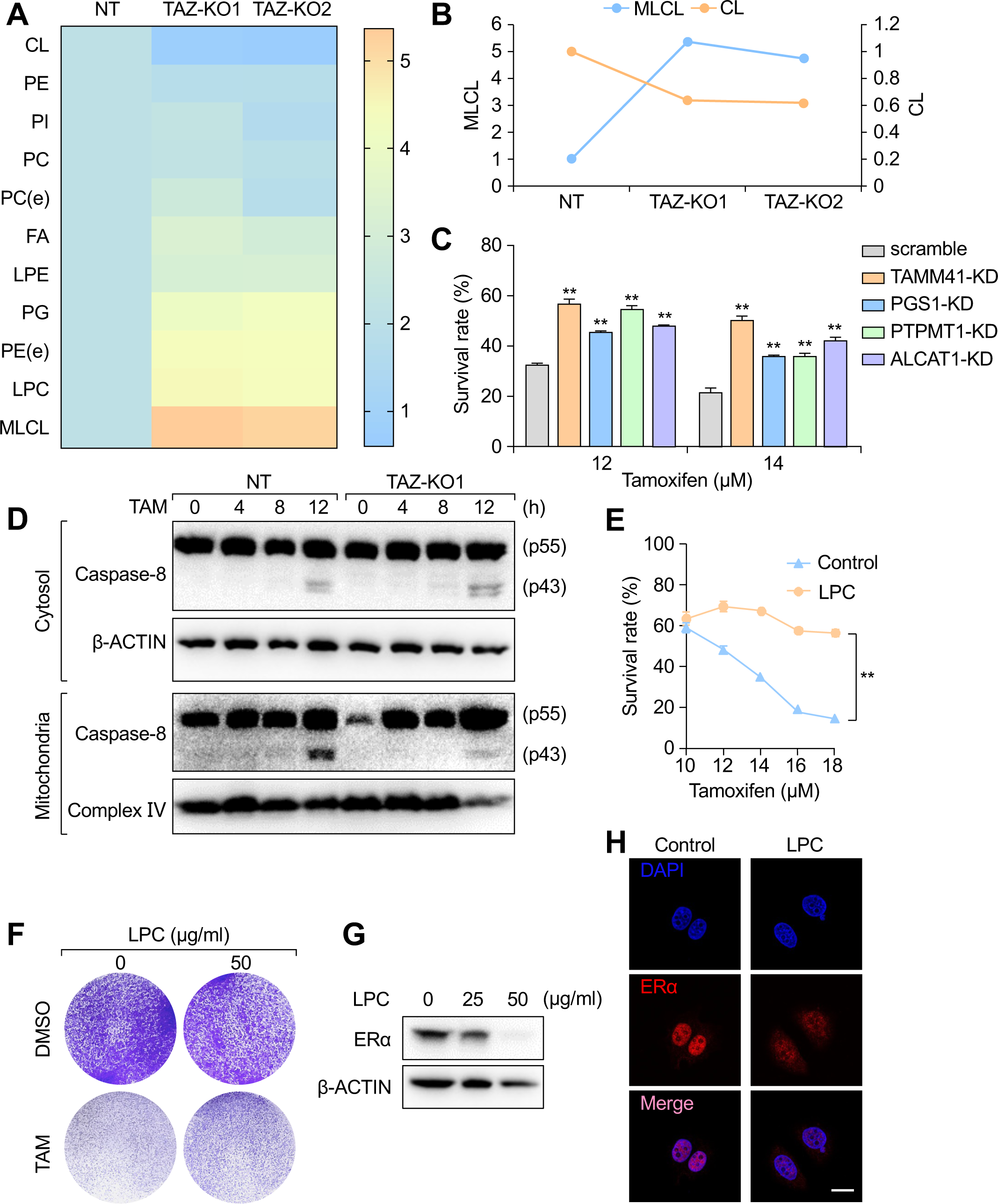
TAZ KO alters cellular phospholipid metabolism and induces tamoxifen resistance. **A.** Phospholipid metabolites in TAZ-KO MCF7 cells were profiled using HPLC/MS. An NT-control and two KO MCF7 stable cell lines were used in the experiment. The level of each phospholipid in NT cells was defined as the base line for plotting. The heat map shows the changes of major phospholipid types in TAZ knockout cells. **B.** The levels of MLCL and mature CL were changed differentially after TAZ KO in MCF7 cells. **C.** Enzymes involved in CL synthesis and maturation were knocked down by shRNAs. Cells were then treated with two different concentrations of tamoxifen for 48 h. Data represents the cell viability normalized to the corresponding DMSO control group. n = 3, data presented as mean ± SD. * p < 0.05, ** p < 0.01 by Student’s t-test. **D.** NT control and TAZ-KO MCF7 cells were treated with tamoxifen with indicated time, and cells were fractionated to cytosolic- and mitochondria-enriched fractions. The status of caspase-8 cleavage in these samples was analyzed by immunoblotting. β-ACTIN was used as the internal loading control. Complex IV was used as the mitochondrial-specific marker and loading control. **E.** MCF7 cells culture was supplemented with vehicle control or LPC. Cells were then treated with DMSO or increasing concentrations of tamoxifen. The cell survival rate for each group was normalized to DMSO control group. n = 3, data presented as mean ± SD. ** p < 0.01 by t-test. **F.** MCF7 cell culture supplemented with vehicle control or LPC were treated with DMSO or with 14 µM tamoxifen. Seven days later, survived cell colonies were stained with crystal violet. n = 3 per group. **G.** The protein expression of ERα in MCF7 cells treated with indicated concentrations of LPC were detected by immunoblotting. β-ACTIN was used as the internal loading control. **H.** MCF7 cells were treated with vehicle control or 50 μg/ml LPC before fixed for immunofluorescent staining against ERα (red). Nuclei were stained with DAPI (blue). Scale bar, 20 μm. **See also Supplemental Figure S5.**

### TAZ-KO increases the cellular LPC level, which is partially responsible for tamoxifen resistance

In the cellular phospholipid network, changes in one species can impact other phospholipids, disrupting homeostasis [45]. Additionally, a previous study showed that TAZ depletion not only decreased cardiolipin but also increased intracellular phosphatidylserine levels, affecting the stemness and differentiation of acute myeloid leukemia cells [24]. Thus, we re-examined the global levels of intracellular phospholipids in TAZ-KO cells and observed an increase in some phospholipids, and lysophosphatidylcholine (LPC) was the most up-regulated lipid besides MLCL (Fig. 5A). To investigate the relationship between elevated cellular level LPC and tamoxifen resistance, we supplemented cell culture with external LPC and tested their ability to induce tamoxifen resistance. We found that the addition of LPC significantly improved cell viability and clone formation in tamoxifen killing assays in both MCF7 and T47D cells (Figs. 5E&F, and Supplementary Fig. S5D). Further, we observed that the addition of LPC restrained ERα expression and impaired the nuclear localization of ERα, similar to the phenotype observed in TAZ-KO cells (Figs. 5G&H). In conclusion, TAZ deficiency leads to elevated intracellular LPC level, which may contribute to tamoxifen resistance, likely by regulating ERα expression and cellular localization.

### LPC-LPA-LPAR axis supports tamoxifen-resistant cells to survive

Excessive LPC can undergo further hydrolysis extracellularly by lysophospholipase D (lysoPLD), such as autotaxin (ATX), resulting in the generation of lysophosphatidic acid (LPA) [46, 47]. LPA is a bioactive molecule that can bind to its receptors (LPAR1-6) and activate intracellular survival signals, promoting tumor cell division, growth, and migration (Fig. 6A) [48]. As the level of LPC is significantly increased in TAZ-deficient tumor cells, we investigated whether the level of LPA is similarly affected and increased. Our results showed that the level of LPA was significantly elevated in the conditional media from TAZ-KO MCF7 cells, with a 1.5-fold upregulation compared to control cells (Fig. 6B). Treatment of TAZ-KO MCF7 cells with the LPAR1/2/3 antagonist, Ki16425, showed that these cells were more sensitive than control cells (Supplementary Fig. S6A). TAZ-deficient MCF7 cells were also re-sensitized to tamoxifen treatment (Fig. 6C). Similar experimental results were obtained in T47D cells (Supplementary Fig. S6B). Besides, external supplement of LPA to cell culture also improved cell viability of MCF7 cells during tamoxifen killing assay (Supplementary Fig. S6C). As TAZ-KO cells down-regulate their ERα signaling to resist tamoxifen therapy, they may simultaneously require the up-regulation of LPA signaling to provide a cellular survival signal. Subsequently, we found that the mRNA level of *ENPP2*, which encodes ATX, was upregulated in TAZ-deficient MCF7 cells (Fig. 6D), and the protein level of ATX in the supernatant and cytosol was significantly elevated compared to control cells (Fig. 6E). In both MCF7 and T47D cells, treatment with the ATX inhibitor PF8380 in combination with tamoxifen lowered the survival rate of TAZ-KO cells to a level comparable to that of control cells (Fig. 6F and Supplementary Fig. S6D). Taken together, these results suggest that the survival of tamoxifen-resistant TAZ-KO cells might depend on the LPC-LPA-LPAR phospholipid signaling axis. Inhibition of this synthesis pathway may restore the sensitivity of ER+ tumor cells to tamoxifen treatment.

**Figure 6.**
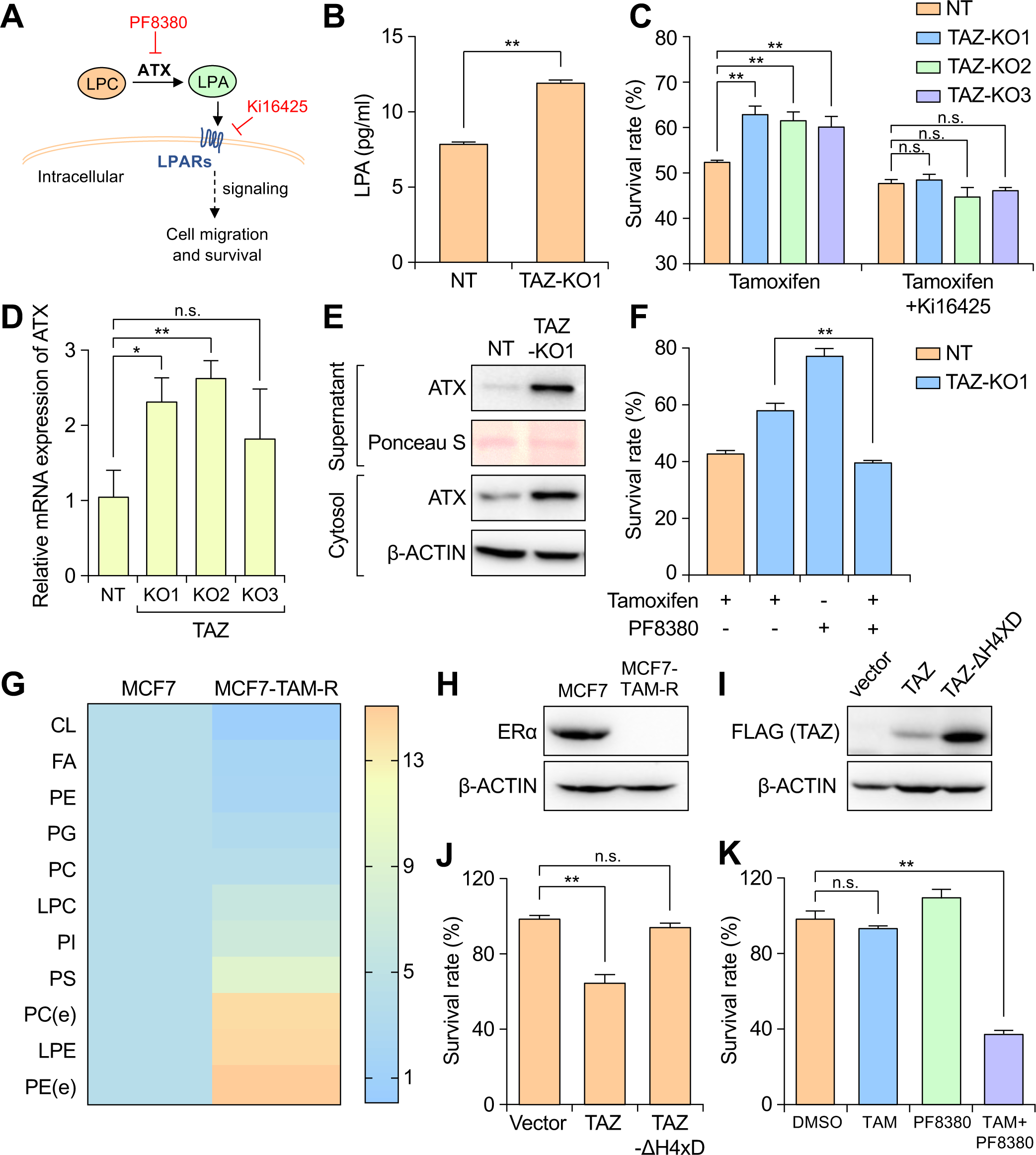
Blockade of the LPA synthesis and signaling pathway re-sensitizes tumor cells to tamoxifen treatment. **A.** A Schematic illustration of the LPC-ATX-LPA axis. **B.** The concentration of LPA in conditioned media from either NT control or TAZ-KO MCF7 cells was determined by ELISA. n = 3, data presented as mean ± SD. ** p < 0.01, by Student’s t-test. **C.** NT control and TAZ-KO MCF7 cells were treated with 10 μM tamoxifen alone or in combination with 20 μM Ki16425 for 48 h. The cell survival rate for each group was presented. n = 3 per group, data presented as mean ± SD. n.s. represents no significant difference, ** p < 0.01 by Student’s t-test. **D.** Relative mRNA expression levels of *ATX* in NT control and in three independent TAZ-KO MCF7 cells were detected by qPCR. *GAPDH* was used as the internal control. n = 3, data presented as mean ± SD. n.s. represents no significant difference, * p < 0.05, ** p < 0.01 by Student’s t-test. **E.** The ATX protein expression levels in conditioned media or in cytosol of NT control and TAZ-KO MCF7 cells were detected by immunoblotting. Ponceau S and β-ACTIN were used as the loading controls. **F.** NT control or TAZ-KO MCF7 cells were treated with either 10 μM tamoxifen alone, 20 μM PF8380 alone, or both tamoxifen and PF8380 for 36 h. The cell survival rate for each group was normalized to DMSO control group. n = 3, data presented as mean ± SD. ** p < 0.01 by Student’s t-test. **G.** Phospholipid metabolites in normal MCF7 cells and in tamoxifen-resistant TAM-R-MCF7 cells were profiled using HPLC/MS. The level of each phospholipid in normal MCF7 cells was defined as the base line for plotting. The heat map shows the changes of major phospholipids in TAM-R-MCF7 cells. **H.** The expression levels of ERα in MCF7 and TAM-R-MCF7 cells were detected by immunoblotting. β-ACTIN was used as the internal loading control. **I.** TAM-R-MCF7 cells were transduced with lentiviruses containing either vector-control, TAZ overexpression cDNA, or TAZ-ΔH4XD mutant construct. The protein expression level of exogenous TAZ was determined by immunoblotting. β-ACTIN was used as the internal loading control. **J.** Stable cell lines generated in experiment in I were treated with 10 μM tamoxifen for 36 h. The cell survival rate for each group was presented. n = 3 per group, data presented as mean ± SD. n.s. represents no significant difference, * p < 0.05, ** p < 0.01 by Student’s t-test. **K.** TAM-R-MCF7 cells were treated with 10 μM tamoxifen alone, 20 μM PF8380 alone, or in combination with both tamoxifen and PF8380 for 36 h. The cell survival rate for each group was presented. n = 3 per group, data presented as mean ± SD. n.s. represents no significant difference, ** p < 0.01 by Student’s t-test. **See also Supplemental Figure S6.**

To extend our findings, we utilized a tamoxifen-resistant MCF7 cell line (MCF7-TAM-R), which was generated through a long-term, low concentration tamoxifen treatment *in vitro*. Using HPLC-MS analysis, we measured the phospholipid metabolites in TAM-R cells and found that MCF7-TAM-R showed similar phospholipid production compared to TAZ-KO cells, with about an 80% decrease in CL and a 2-fold increase in LPC (Fig. 6G). Furthermore, the protein expression level of ERα in MCF7-TAM-R was significantly low, similar to TAZ-KO cells (Fig. 6H). The overexpression of TAZ, but not TAZ-ΔH4XD mutant, restored tamoxifen sensitivity of MCF7-TAM-R cells (Figs. 6I&J). Additionally, we employed the ATX inhibitor, PF8380 to treat the resistant cells with or without tamoxifen. Neither tamoxifen nor PF8380 alone had a significant killing effect on MCF7-TAM-R cells. However, the combination of both PF8380 and tamoxifen significantly reduced the survival rate of tumor cells by nearly 70% (Fig. 6K).

## Discussion

The occurrence of therapy resistance in breast cancer is a severe bottleneck in clinical and is the major cause of treatment failure and patient mortality [2, 4, 49-51], and the molecular mechanisms that underlie therapy resistance are not yet fully understood. To address this challenge, we employed a genome-scale CRISPR-Cas9 loss-of-function screen to systematically identify candidate therapy-resistance genes. Our study revealed that TAZ, a mitochondrial-localized lysophospholipid-phospholipid transacylase that is required for the production of the phospholipid, CL, is essential for tamoxifen-induced cell death [23].

Lack of expression of the estrogen receptor alpha (ERα), the cellular target of tamoxifen, is a well-known mechanism of tamoxifen resistance in many studies [52, 53]. Our results showed that loss of TAZ led to suppression of ERα expression, retention of ERα in the cytoplasm, and perturbation of ER downstream signaling, including CCND1, which is essential for G1/S cell cycle transition. Therefore, TAZ appears to downregulate ERα, the major tamoxifen target, as a means of escaping tamoxifen therapy. TAZ-KO cells exhibited a moderate reduction in cell cycle progression rate due to the downregulation of ER signaling. Although cell cycle inhibition is a primary strategy for controlling tumor growth and promoting apoptosis [54], quiescent cells or dormancy tumor cells also proliferate slower or even halt their cell cycle [39, 55-57]. In addition, drug-tolerant persister (DTP) cells have emerged as a concept in drug resistance research, characterized by a quiescent or slowly-cycling state in which tumor cells enter a reversible cell cycle arrest. Several DTP-related studies have been conducted to identify the genetic features or epigenetic modifications that regulate the development of DTP-induced drug resistance [40, 41, 58, 59]. We knocked out a set of genes with low-, medium-, or normal (high)-proliferation indices and examined their tamoxifen resistance. KO of slow-proliferating genes significantly increased tamoxifen resistance, while KO of the remaining genes had little effect. Thus, cell cycle regulation can be a “double-edged sword” for tumor therapeutic resistance because slow-cycling cells are more likely to be drug-resistant.

We discovered that TAZ deficiency leads to altered ERα signaling and tamoxifen resistance through its metabolic enzyme activity. Firstly, we observed a reduction in mature CL levels in TAZ knockout ER+ cells, and CL was found to be required for maintaining tamoxifen sensitivity. A previous study demonstrated that CL is necessary for caspase-8 activation induced by exogenous apoptosis (33). Tamoxifen promotes apoptosis via a caspase-8 dependent mechanism as a calmodulin antagonist [60, 61], and our results show that tamoxifen induces apoptosis of breast cancer cells by activating caspase-8. Abnormal CL metabolism caused by TAZ deficiency inhibits the apoptotic response. Secondly, lipid metabolism is a highly interconnected network, and alteration of one lipid species can affect other intracellular lipids [45]. Depletion of TAZ blocks the remodeling of mature CL from MCLC, resulting in the accumulation of MLCL and other upstream phospholipids in tumor cells. As an indirect result of the perturbation of the CL synthesis pathway, many other phospholipids, including LPC, are upregulated in TAZ-KO cells. Our results show that LPC induces tumor cells to develop tamoxifen therapy resistance, causing the down-regulation of the ERα protein and reducing its nuclear localization. Thus, the down-regulation of ERα signaling caused by TAZ KO could be mainly mediated through LPC.

The LPC generated as a result of TAZ deficiency could be further metabolized into bioactive LPA by the enzyme ATX, whose expression is known to be associated with tumor progression [47, 62, 63]. A few studies report that increased ATX expression and LPA signaling promote the development of cancer resistance to chemotherapy, radiotherapy, and immunotherapy [48, 64, 65]. However, the relationship between the ATX-LPA axis and tamoxifen resistance is not well understood. In our study, we found that inhibiting ATX enzymatic activity directly or blocking LPA receptor activation sensitizes tumor cells to tamoxifen treatment. The ATX-LPA axis has been implicated in many other diseases, including pulmonary fibrosis, atherosclerosis, inflammatory arthritis. Some ATX inhibitors have been developed and entered into clinical trials, including GLPG1690, BBT-877 and BLD-0409[66]. Currently, several small-molecule ATX inhibitors are being tested in human clinical trials for pulmonary fibrosis [67]. These inhibitors could be repurposed for usage in treating tamoxifen-resistant breast cancer patients.

## Materials and Methods

### Cell culture

HEK293T, MCF7, SCP2, and MDA-MB-231 cell lines were cultured in Dulbecco’s modified Eagle’s medium (DMEM, Gibco, Catalog # C11995500BT) supplemented with 10% heat-inactivated Fetal Bovine Serum (FBS, Gemini, Catalog # 900-108), 1% penicillin and streptomycin (P/S, Gibco, Catalog # 15140163). T47D and ZR-75-1 cell lines were cultured in 1640 medium (Gibco, Catalog # C11875500CP) supplemented with 10% FBS and 1% P/S. The tamoxifen-resistant (TAM-R) MCF7 cell strain was kindly provided by Prof. Z. Chang in Tsinghua University. Briefly, this cell strain was generated by culturing MCF7 cells with low-dose tamoxifen for an extended time before gradually increasing the tamoxifen concentration in the culture media until populations of cells could grow normally in this high tamoxifen culture condition. All cell lines were cultured in a constant temperature incubator at 37 °C and 5% CO_2_. The cell lines were purchased from National Collection of Authenticated Cell Cultures in China. and tested monthly for mycoplasma contamination.

### CRISPR-Cas9 screen

The Human Brunello v2 CRISPR KO pooled library (76,441 gRNAs, targeting 19,114 genes, including 1,000 controls) was bought from Addgene (Watertown, MA, USA, Addgene ID: 73179). Plasmid library were amplified by electro-transformation (Lucigen, Catalog # 60242-1). Lentivirus library was packaged by co-transfecting HEK293T with psPAX2 and pMD2.G plasmids, using PEI transfection method (SIGMA, Catalog # 408727). The viruses were collected at 48 h and 72 h after transfection. Estrogen receptor-positive breast cancer cell line MCF7 was infected by the above virus at an MOI of 0.2. Transduced cells were selected with 3 μg/ml puromycin (Solarbio, Catalog # P8230) for 3 days. Library cells were treated with either 12 μM tamoxifen (Abacm, Catalog # ab120656) or DMSO (Solarbio, Catalog # D8371) for 72 h. During the screen, cells were passaged maintaining a minimum coverage of 500 cells per sgRNA. Cells were collected and extracted for genomic DNA. Then, sgRNA regions were amplified using staggered primers (see below) by Ex Taq (TaKaRa, Catalog # RR001B) to construct a library for next-generation sequencing. The PCR products were extracted by a gel extraction kit (TIANGEN, Catalog # DP209-03). Data analysis was performed using MAGeCK.

The P5 primer: 5’AATGATACGGCGACCACCGAGATCTACACTCTTTCCCTACACGACGCTCTTCCGATCT[s]T TGTGGAAAGGACGAAACACCG 3’

The P7 primer:

5’CAAGCAGAAGACGGCATACGAGATNNNNNNNNGTGACTGGAGTTCAGACGTGTGCTCTTCCGATCTCCAATTCCCACTCCTTTCAAGACCT 3’, NNNNNNNN is the barcode region. The sgRNAs used in candidate gene confirmation were listed in Supplementary Table S1.

### TIDE analysis

Genomic DNA was purified from the indicated cells and the sgRNA-targeted loci were amplified by PCR (TSINGKE, Catalog # TSE101). The primers designed for PCR are 200-250 bp upstream and about 250 bp downstream from the sgRNA-targeted loci. KO efficiency was analyzed by the indel efficiency at https://tide.nki.nl.

### Drug killing assay

Cells were seeded in 12-well plates (5×10^4^ for MCF7, SCP2, or MDA-MB-231 cells per well, 8×10^4^ for T47D, or ZR75-1 cells per well). After about 24 h, cells were treated with different agents (Tamoxifen citrate, Abcam, Catalog # ab120656; Cisplatin, Harvey, Catalog # HZB0054; Olaparib, Selleck, Catalog # S1060). Tamoxifen citrate is reported to have the IC50 in MCF7 of 10.2 μM[68]. The concentrations used in this study is on par with the IC50 concentration of tamoxifen citrate. In the experimental end point, 200 μl luciferase lysis buffer (2 mM EDTA, 20 mM DTT, 10% glycerol, 1% Triton X-100, 25 mM Tris-base, pH 7.8) was added into each well. Cells were lysed at room temperature for one hour. 25 μl cell lysate was transferred into a 96-well white plate (triplicates for each sample), then 75 μl luciferase substrate solution (25 mM glycylglycine, 15 mM potassium phosphate, 4 mM EGTA, 2 mM ATP, 10 mM DTT, 1 mM D-luciferin, 15 mM MgSO4, pH 7.8) was added into each well. Luciferase activity was measured by EnVision® Multimode Plate Reader (PerkinElmer).

### Correlation analysis of cell proliferation and resistance

2×10^4^ NT control or specific gene KO MCF7 cells were seeded in 12-well plates, cultured for 72 h, and then treated with DMSO or tamoxifen for another 72 h. Cells were then lysed for luciferase assay to determine their relative cell number in each group. The “proliferation index” for each gene was calculated as the relative cell number of KO group divided by the relative cell number of NT control group in DMSO treatment. The “resistance index” for each gene was calculated as the relative cell viability of KO group divided by the relative cell viability of NT control group in tamoxifen treatment.

### Colony formation assay

1x10^4^ MCF7 cells were seeded per well. The cell culture medium was replenished every 3 days. The colonies were cultured for 1 week with or without tamoxifen, then fixed with 4% paraformaldehyde (Biosharp, Catalog # BL539A), and stained by crystal violet (Solarbio, Catalog # C8470).

### Tumor sphere formation

The 12-well plates used for tumor sphere formation were pre-treated with 12 mg/ml poly HEMA (SIGMA, Catalog # P3932) and dried on a clean bench. MCF7 cells used for tumor sphere formation were trypsinized to generate single-cell suspension. 5 x10^3^ MCF7 were plated to 1 ml DMEM/F12 (Invitrogen, Catalog # C11765500BT) medium containing B27 (Invitrogen, Catalog # 17504044), 20 ng/ml bFGF (Novoprotein, Catalog # C046), 20 ng/ml EGF (Macgene, Catalog # CC102), penicillin/streptomycin. Fresh medium was added against the sides of the well every 3 days. The tumor spheres were generated after about 7 days.

### Reverse transcription and qPCR

The total RNA of cells was isolated using RNAiso Plus Regent (Takara, Catalog # 9108) following the manufacturer’s instructions. RNA was reverse transcribed into cDNA using ReverTra Ace® qPCR RT Master Mix (TOYOBO, Catalog # CC102FSQ-201). Quantitative PCR (qPCR) was performed by ABI Quant Studio 3 series PCR machine (Applied Biosystem) using the Power Green qPCR Mix (DONGSHENG BIOTECH, Catalog # CC102P2105). The gene-specific primer sets were used at a final concentration of 0.5 μM. All qPCR assays were performed in triplicates. Relative mRNA expression values of each target gene were normalized to *GAPDH*. The qPCR primer sequences used for examined genes were listed in Supplementary Table S3.

### Virus production and stable cell line generation

Lentiviruses were produced by co-transfecting HEK293T cells with the lentiviral vectors and packaging plasmids, psPAX2 and pMD2.G, using PEI (SIGMA, Catalog # 408727) according to the manufacturer’s recommendations. Viral supernatants were collected at 48 h and 72 h post-transfection, filtered through 0.45 μm PVDF syringe filters (Millex, Catalog # SLHV033RB). Viral transductions were performed for about 12-18 hours in the presence of 8 μg/ml polybrene (Macgene, Catalog # MC032). Cells were expanded for another 3-4 days for FACS sorting or antibiotic selection at appropriate concentrations. Stable overexpression of all genes was achieved using the pLEX-MCS or pLVX-CMV-MCS-mPGK lentiviral plasmids. Stable Knockout of all genes was achieved using lentiCRISPR v2 plasmid. Stable knockdown of all genes was achieved using pLKO.1 lentiviral plasmid. shRNA sequences used to knockout related genes were listed in Supplementary Table S4.

### Immunoblotting analysis

Whole cell lysate samples were collected using cell lysis buffer (50 mM Tris-HCl pH 7.4, 150 mM NaCl, 1 mM EDTA, and 1% NP-40, add protease inhibitor cocktail when use) on ice. Lysates were heated at 95 °C for 10 min to denature the proteins, loaded to SDS-PAGE gel for electrophoresis, and subsequently transferred to PVDF membranes (Millipore, Catalog # IPVH00010). Membranes were blocked in 5% skimmed milk for 1 h at room temperature before overnight incubation with primary antibodies. Primary antibodies used were: anti-β-ACTIN (1:5,000, Abcam, Catalog # ab6276), anti-FLAG (1:2,000, CST, Catalog # 14793), anti-HA (1:2,000, Roche, Catalog # 11867423001), anti-ERα (1:2,000, CST, Catalog # 8644), anti-Tafazzin (1:100, Santa Cruz, Catalog # sc-365810), anti-GREB1 (1:1,000, Proteintech, Catalog # 28699-1-AP), anti-Cyclin D1 (1:1,000, Immunoway, Catalog # YT1173), anti-TFF1 (1:1,000, Abcam, Catalog # ab92377), anti-caspase-8 (1:1,000, CST, Catalog # 9746T), anti-Complex IV (1:2,000, Proteintech, Catalog # 11242-1-AP), anti-ATX (1:2,000, Immunoway, Catalog # YT5359). Membranes were incubated with secondary antibodies for 1 h at room temperature. Secondary antibodies used were: Goat Anti-Mouse IgG (H&L)-HRP conjugated (1:5,000, Easybio, Catalog # BE0102), Goat Anti-Rabbit IgG (H&L)-HRP conjugated (1:5,000, Easybio, Catalog # BE0101), Goat Anti-Rat IgG (H&L)-HRP Conjugated (1:5,000, Easybio, Catalog # BE0108). The chemiluminescence signals were detected by ECL substrate (Tanon, Catalog # 180-5001) on Champ Chemi Digital Image Acquisition Machine (Beijing Sage Creation Science Co, LTD). Image quantifications were performed by ImageJ (NIH).

### Immunoprecipitation

The overexpression plasmids as indicated were transfected into HEK293T cells in six-well plates using PEI. After 24-48 h, cell lysates were harvested and immunoprecipitated with 5 μl of magnetic anti-FLAG-beads (Sigma, Catalog # M8823) each sample overnight at 4 °C with rocking. After extensive washing (three to five times) with lysis buffer containing protease inhibitor cocktail (Cwbio, Catalog # CW2200S), the beads were spun down and resuspended with 50 μl loading buffer. After boiling for 10 min at 95 °C, protein samples were loaded to SDS-PAGE gel along with 10% input samples for electrophoresis and subsequently transferred to PVDF membranes, and then detected with appropriate antibodies.

### Immunofluorescence staining

Cells were cultured on coverslips in a 12-well plate. The medium was discarded and the coverslips were washed twice with cold PBS (Solarbio, Catalog # P1020). Cells were then fixed with 4% paraformaldehyde for 15 minutes at room temperature, washed with PBS 3 times, blocked and permeabilized in PBS containing 5% goat serum (Solarbio, Catalog # SL038) and 0.2% Triton X-100 (Solarbio, Catalog # T8200) for one hour at room temperature. Cells were then stained with respective primary antibodies overnight at 4 °C. Fluorophore labeled secondary antibodies incubated cells for 1 or 2 hours at room temperature in dark. The primary antibodies or fluorescent probe used were anti-FLAG (1:200, CST, Catalog # 14793), anti-ERα (1:200, CST, Catalog # 13258), and Mito-Tracker Red CMXRos (Beyotime, Catalog # C1035). Secondary antibodies used in the were Dylight649 Goat Anti-Rabbit IgG(H+L) (1:100, EarthOx, Catalog # E032620). Images were captured by Olympus FV3000 confocal microscope.

### Lipid extraction

10^6^-10^7^ cells were used to extract lipids for HPLC/MS. Cells were washed with cold PBS three times. Samples were resuspended in 2 ml PBS, and then added to 2:1 chloroform: methanol (v/v) on ice (Chloroform, J.T.Baker, Catalog # 67-66-3; Methanol, Fisher, Catalog # 67-56-1). Samples were vortexed for 1 min and stand for 10 min, repeated three times. Samples were centrifuged for 15 minutes at 1,000 g. The organic layer was collected and the methanol/water layer was discarded. Samples were stored in chloroform at 4 °C until further analysis.

### Cell cycle analysis

Cells were collected, washed by PBS once, and fixed in 70% cold ethanol overnight at 4 °C. Samples were washed by PBS again, treated with 20 μg/ml RNase A (Beyotime, Catalog # ST576) for 30 minutes at 37 °C, then stained with propidium iodide (PI, Biolegend, Catalog # 421301) for 30 minutes at 4 °C. Protected from light before being subject to FACS analysis. Calibur (BD) flow cytometry was used. Data analysis was performed on Modfit or Flow Jo software.

### Xenograft experiments

All procedures involving mice and experimental protocols were approved by the Institutional Animal Care and Use Committee (IACUC) of Tsinghua University. Nude/nude mice were purchased from Vital-River. All mice were maintained under specific pathogen-free conditions. Self-made slow-release oestrogen pellets (β-oestradiol 0.72 mg per implantation) were transplanted into 6-week-old female Nude/nude mice before tumor cell inoculation. 50 μl PBS containing 2×10^6^ TAZ KO MCF7 or control cells (NT) were mixed 1:1 (volume) with Matrigel (Corning, Catalog # 354234) and then injected into the fourth mammary fat pads of mice. When the tumor size reached about 60 mm^3^, mice were randomized to be treated with tamoxifen (Abcam, Catalog # ab120656) or vehicle (10% DMSO, 90% corn oil). Tamoxifen was injected intraperitoneally at a dose of 1.5 mg per mouse, three times per week. Tumor volumes were measured every four days and the tumor volumes were calculated as volume (mm^3^) = length × width^2^/2. No blinding was done.

### ELISA

The lysophosphatidic acid (LPA) concentration in cell supernatant was measured by ELISA (Abmart, Catalog # AB-3859A) according to the manufacturer’s protocol. Cells were cultured in serum-free medium for 18 h. The supernatant was collected and centrifuged at 3000 g for 10 minutes to remove debris. Plates were measured by EnVision® Multimode Plate Reader (PerkinElmer).

### Cellular fractionation

Cells were washed by PBS, pelleted and lysed in cytosol extraction buffer (50 mM HEPES pH 7.4, 150 mM NaCl, 1M hexylene glycol, 100 μM digitonin) for 10 minutes on ice. The lysates were centrifuged at 500 g for 5 minutes at 4 °C and the supernatants were cytosolic fraction. Then the pellets were further lysed in membrane extraction buffer (50 mM HEPES pH 7.4, 150 mM NaCl, 1M hexylene glycol, 1% IGEPAL) for 10 minutes on ice. The lysates were centrifuged at 3,000 g for 5 minutes at 4 °C and the supernatants were membrane fraction. The remaining pellets were nuclei fraction.

### Quantification and Statistical Analysis

Sample sizes were chosen based on preliminary experiments or experimental designs in related studies. For animal studies, no statistical methods were used to estimate sample size. Results were reported as mean ± SD (standard deviation) or mean ± SEM (standard error of the mean), as indicated in the figure legends. Unpaired t-test or two-way ANOVA analysis was used to calculate the statistical significance of the difference in a particular measurement between groups. p value less than 0.05 was considered to be statistically significant. All experiments with representative images (including immunoblotting and immunofluorescence staining) were repeated at least twice and representative images are shown.

## Acknowledgments

We thank D. Pan for helping with MAGeCK analysis; Z. Chang and Q. Ye for generously sharing the tamoxifen resistance cell; We thank all members of the Zheng laboratory for helpful discussions and technical assistance. We thank the Technology Center for Protein Sciences at Tsinghua University for mass-spectrometry support, the Laboratory Animal Research Center for animal research support, and the Center of Biomedical Analysis at Tsinghua University for FACS and optical imaging support. The study was partially supported by the National Key Research and Development Program of China (2020YFA0509400 to H.Z.), the National Science Foundation of China (81772981 and 81972462 to H.Z.), the Tsinghua University Initiative Scientific Research Program, and the Tsinghua-Peking Center for Life Sciences.

## Author contributions

X.L. and H.Z. designed the overall study. X.L. performed the experiments, analyzed data, and wrote the manuscript. T.Z. provided technical help for next-generation data analysis. L.Z. provided help with the CRISPR/Cas9 screening. Y.Z. provided technical help for animal work. H.H. provided technical help for cell works. H.Z. supervised the overall study. H.Z., C.L., and X.L. revised the manuscript.

