## Supplementary information for "Tafazzin Mediates Tamoxifen Resistance by Regulating Cellular Phospholipid Composition in ER-Positive Breast Cancer"

**SUPPLEMENTAL INFORMATION**

**Supplementary Data**

Supplementary Fig. 1 is related to Fig. 1.

Supplementary Fig. 2 is related to Fig. 2.

Supplementary Fig. 3 is related to Fig. 3.

Supplementary Fig. 4 is related to Fig. 4.

Supplementary Fig. 5 is related to Fig. 5.

Supplementary Fig. 6 is related to Fig. 6.

**Figure S1. The expression and survival analysis of candidate genes in patient datasets, related to Figure 1. A.** The initial quality check of sgRNA library was performed by analyzing the distribution of sgRNAs in sgRNA library transduced cell population, which was used in later screening. **B.** Top candidate genes for enriched sgRNAs. Candidates were plotted based on mean log2 fold change and P values computed by MaGeCK. **C.** Top candidate genes for depleted sgRNAs. Candidates were plotted based on mean log2 fold change and P values computed by MaGeCK. **D.** Several candidate genes were selected and the stable KO of these genes were generated in MCF7 cells. Cells were treated with DMSO or 12 µM tamoxifen for 48 h. The cell survival rate for each group was presented. n = 3 per group, data presented as mean ± SD. n.s. represents no significant difference, * p < 0.05, ** p < 0.01 by Student’s t-test. **E.** Kaplan-Meier survival plot shows the correlation between the mRNA expression of *ROMO1* and the relapse-free survival (RFS) of ER+ breast cancer patients with tamoxifen therapy based on an online database (<https://kmplot.com>). **F.** Kaplan-Meier survival plot shows the correlation between the mRNA expression of *TAZ* and the relapse-free survival (RFS) of ER+ breast cancer patients based on an online database (https://kmplot.com). **G.** Kaplan-Meier survival plot shows the correlation between the mRNA expression of *PELO1* and the relapse-free survival (RFS) of ER+ breast cancer patients with tamoxifen therapy based on an online database (https://kmplot.com). **H.** The mRNA expression level of *ROMO1* is higher in breast tumor tissues compared to normal tissues in both Luminal A and Luminal B subtypes of breast cancers. Data was analyzed based on TCGA and GTEx data set on GEPIA2 (http://gepia2.cancer-pku.cn). TPM, Transcripts per million.

**Figure S2. Loss-of-TAZ has little effect of therapy resistance in ER-negative breast cancer, related to Figure 2. A-C.** TAZ KO in MCF7 (A), T47D (B) and ZR-75-1 (C) cells using 3 independent sgRNAs were performed and the KO efficiencies were analyzed by TIDE assay. **D.** Representative images of the tumor spheres in Fig. 2D. Scale bar, 50 μm. **E.** Representative images of the tumors in Fig. 2F. **F, G.** TAZ KO in MDA-MB-231 (D) and SCP2 (E) cells using 3 sgRNAs were performed and the KO efficiencies were analyzed by TIDE assay. **H.** NT control and TAZ-KO MDA-MB-231 cells were treated with PBS or increasing concentrations of cisplatin for 24-48 h. The cell survival rate for each group was plotted. n = 3, data presented as mean ± SD. n.s. represents no significant difference by t-test. **I.** NT control and TAZ-KO SCP2 cells were treated with PBS or increasing concentrations of cisplatin for 24-48 h. The cell survival rate for each group was plotted. n = 3, data presented as mean ± SD. n.s. represents no significant difference by t-test.

**Figure S3. Cell cycle-arrest positively correlates with therapy resistance, related to Figure 3. A.** The protein expression levels of TAZ and ERα in NT control and three independent TAZ-KO ZR-75-1 cells were determined by immunoblotting. β-ACTIN was used as the internal loading control. **B.** The expression levels of indicated proteins in NT control and TAZ-KO MCF7 (left) and ZR-75-1 (right) cells were determined by immunoblotting. β-ACTIN was used as the internal loading control. **C.** Cell cycle analysis by PI staining and FACS analysis of NT control and TAZ-KO SCP2 cells. n = 3, data presented as mean ± SD, n.s. represents no significant difference by Student’s t-test. **D, E.** Cell cycle analysis by PI staining and FACS analysis of MCF7 (C) and T47D (D) cells cultured in media containing indicated concentrations of FBS for 24 h**.** n = 3, data presented as mean ± SD, n.s. represents no significant difference, * p < 0.05, ** p < 0.01 by Student’s t-test. **F.** The average cell cycle distribution of NT control, tamoxifen-resistant cells (resistance ratio > 1.5) or non-resistant cells (resistance ratio < 1.5) defined by our screening based on experiment in Fig. 3G.

**Figure S4. TAZ affects tamoxifen resistance through its enzymatic activity, related to Figure 4. A.** HEK293T were transfected with indicated expression vectors. Cells were lysed for immunoprecipitation with α-FLAG antibody for TAZ detection and α-HA antibody for ERα detection. **B, C.** The levels of CL (B) and MLCL (C) were detected using HPLC in TAZ-KO MCF7 cells with either wild type TAZ or TAZ-ΔH4xD mutant rescue. **D.** Endogenous TAZ was KD by shRNA in MCF7 cells, a scramble (Scra) control shRNA was used as control. Wild type TAZ or TAZ-ΔH4xD was re-expressed in either scramble control or TAZ-KD cells. The relative mRNA expression level of *TAZ* was confirmed by qPCR. *GAPDH* was used as the internal control. n = 3, data presented as mean ± SD, ** p < 0.01 by Student’s t-test. **E.** Cell lines generated from experiment in B were treated with DMSO or tamoxifen for 48 h. The cell survival ratio for each group was calculated by normalizing to the survival rate of scramble control group. n = 3, data presented as mean ± SD, ** p < 0.01 by Student’s t-test.

**Figure S5. KO of TAZ alters cellular levels of phospholipids and induces tamoxifen resistance, related to Figure 5.** **A.** Schematic representation of phospholipid metabolism pathway leading to mature CL production, in which TAZ utilizes PC or phosphatidylethanolamine (PE) as acyl chain donors to attach to reacylate monolysocardiolipin (MLCL). **B.** KD of CL metabolism-related enzymes in MCF7 cells. The relative mRNA expression levels of these genes were analyzed by qPCR. *GAPDH* was used as the internal loading control. n = 3, data presented as mean ± SD, ** p < 0.01 by Student’s t-test. **C.** A Schematic illustration of caspase-8 cleavage. **D.** T47D cells were cultured with control vehicle or LPC supplement. Cells were then treated with increasing concentrations of tamoxifen. The cell survival ratio for each group was calculated by normalizing to the survival rate of DMSO-treated control group. n = 3, data presented as mean ± SD. ** p < 0.01 by t-test.

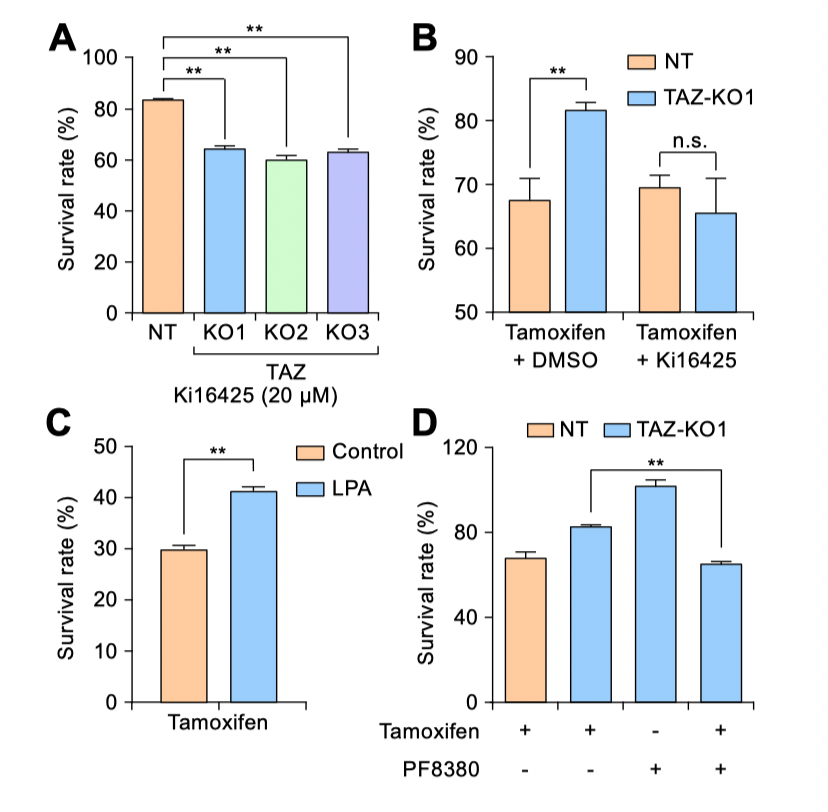

**Figure S6. Blockade of the LPA synthesis axis attenuates tamoxifen resistance, related to Figure 6. A.** NT control or three independent TAZ-KO MCF7 cells were treated with 20 μM Ki16425 for 48 h. The cell survival ratio for each group was calculated by normalizing to DMSO control group. n = 3, data presented as mean ± SD. ** p < 0.01, by Student’s t-test. **B.** NT control or TAZ-KO T47D cells were treated with 10 μM tamoxifen alone or in combination with 20 μM Ki16425 for 48 h. The cell survival ratio for each group was calculated by normalizing to DMSO control group. n = 3, data presented as mean ± SD. n.s. represents no significant difference, ** p < 0.01, by Student’s t-test**. C.** Cell viability of MCF7 under 10 μM tamoxifen treatment, supplemented with vehicle control or LPA. Data presented as mean ± SD. ** p < 0.01 by Student’s t-test. **D.** NT control or TAZ-KO T47D cells were treated with 10 μM tamoxifen alone, 20 μM PF8380 alone, or in combination of tamoxifen and PF8380 for 36 h. The cell survival ratio for each group was calculated by normalizing to DMSO control group. n = 3, data presented as mean ± SD. n.s. represents no significant difference, ** p < 0.01 by Student’s t-test.
