## Supplemental Figures for "Tafazzin Mediates Tamoxifen Resistance by Regulating Cellular Phospholipid Composition in ER-Positive Breast Cancer"

### Supplemental Figure S1

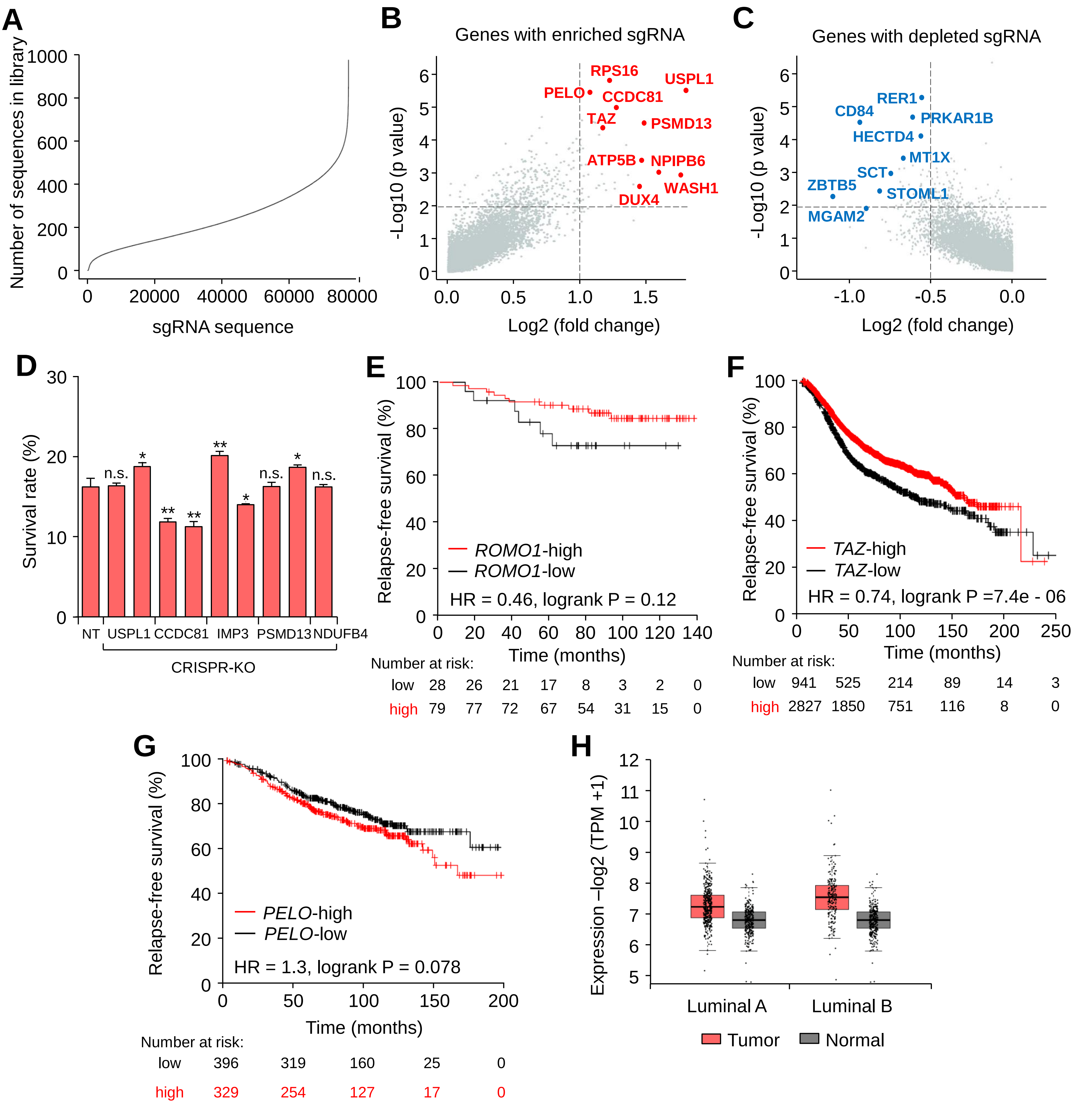

### Supplemental Figure S2

**A**

| MCF7 TIDE analysis | |
|---|---|
| sgRNA | total efficiency (%) |
| 1 | 76.9 |
| 2 | 88.8 |
| 3 | 26.8 |

**B**

| T47D TIDE analysis | |
|---|---|
| sgRNA | total efficiency (%) |
| 1 | 91.7 |
| 2 | 91.4 |
| 3 | 23.6 |

**C**

| ZR-75-1 TIDE analysis | |
|---|---|
| sgRNA | total efficiency (%) |
| 1 | 82.4 |
| 2 | 79.3 |
| 3 | 16.7 |

**D**

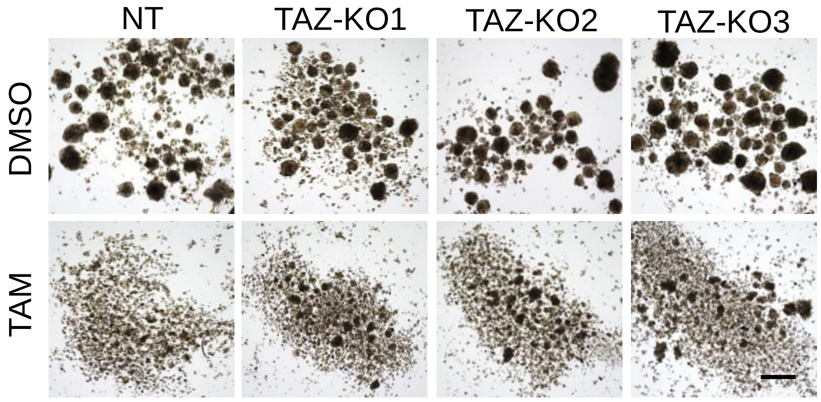


**E**

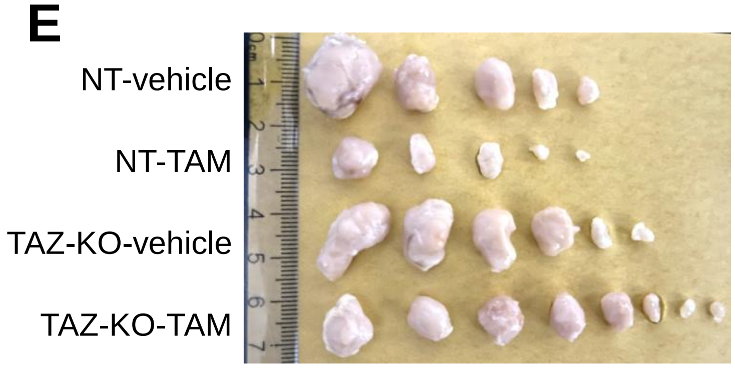


**F**

| MDA-MB-231 TIDE analysis | |
|---|---|
| sgRNA | total efficiency(%) |
| 1 | 89.2 |
| 2 | 91.1 |
| 3 | 66.9 |

**G**

| SCP2 TIDE analysis | |
|---|---|
| sgRNA | total efficiency(%) |
| 1 | 93.2 |
| 2 | 87.7 |
| 3 | 49.6 |

**H**

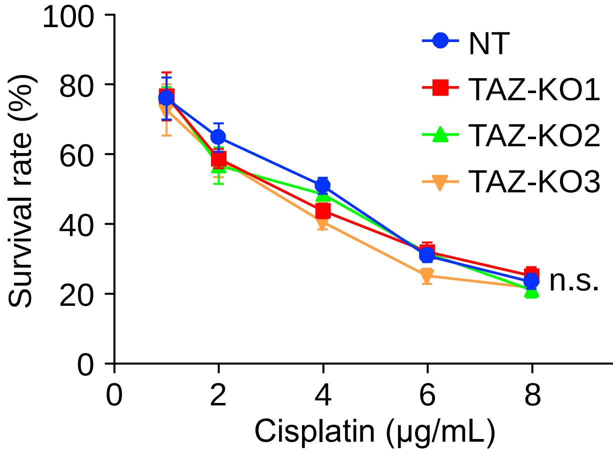


**I**

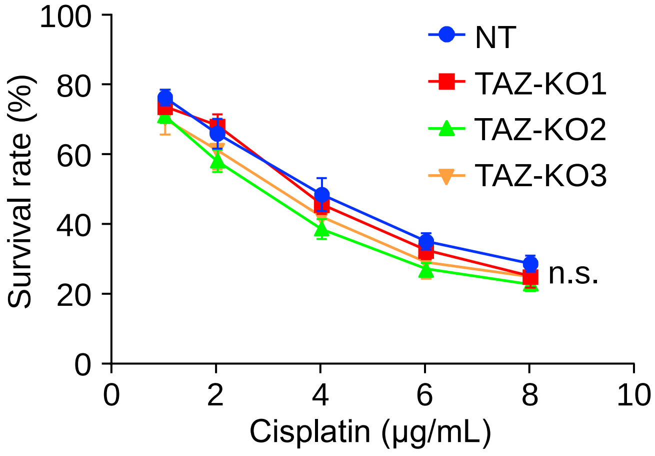

### Supplemental Figure S3

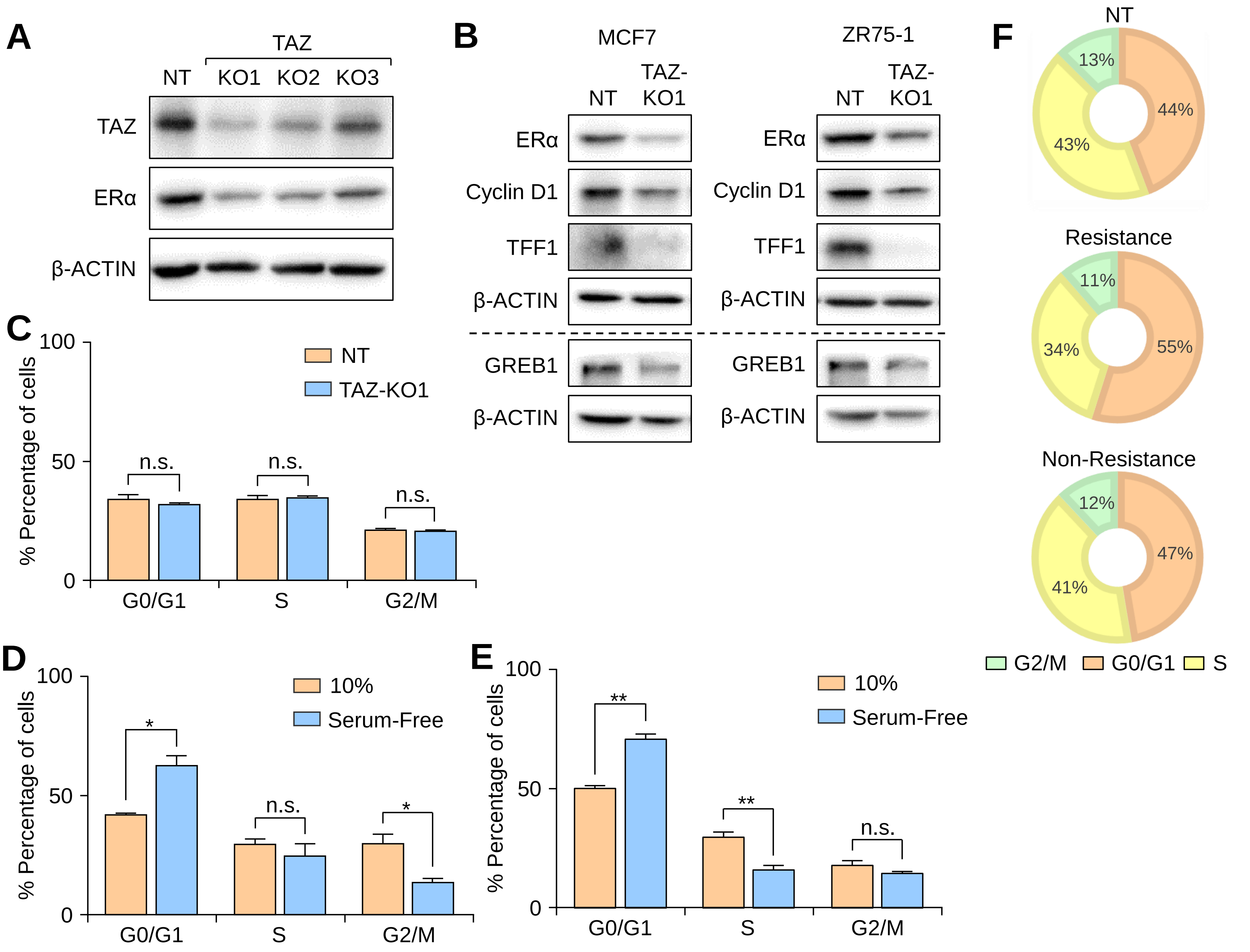

### Supplemental Figure S4

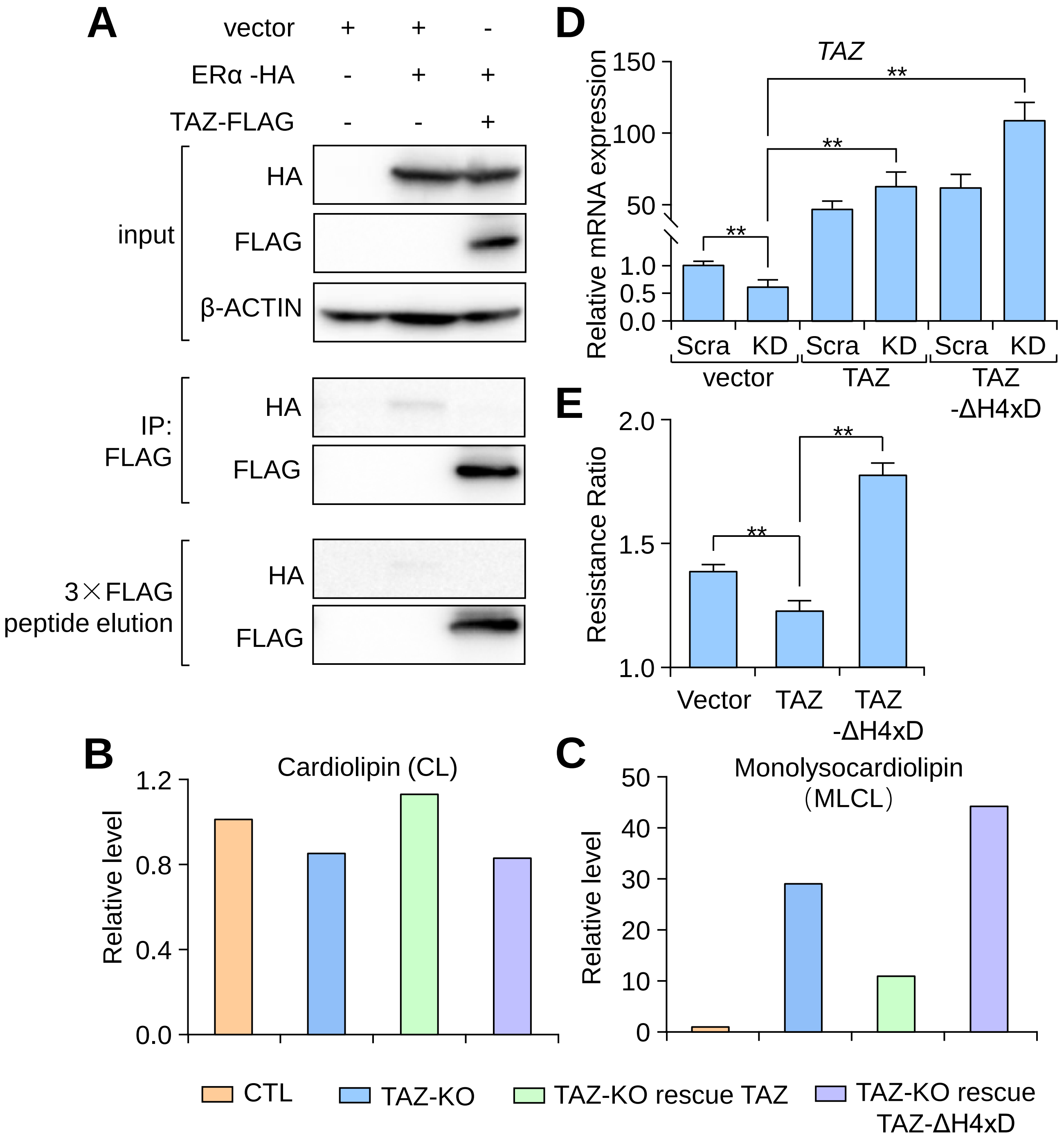

### Supplemental Figure S5

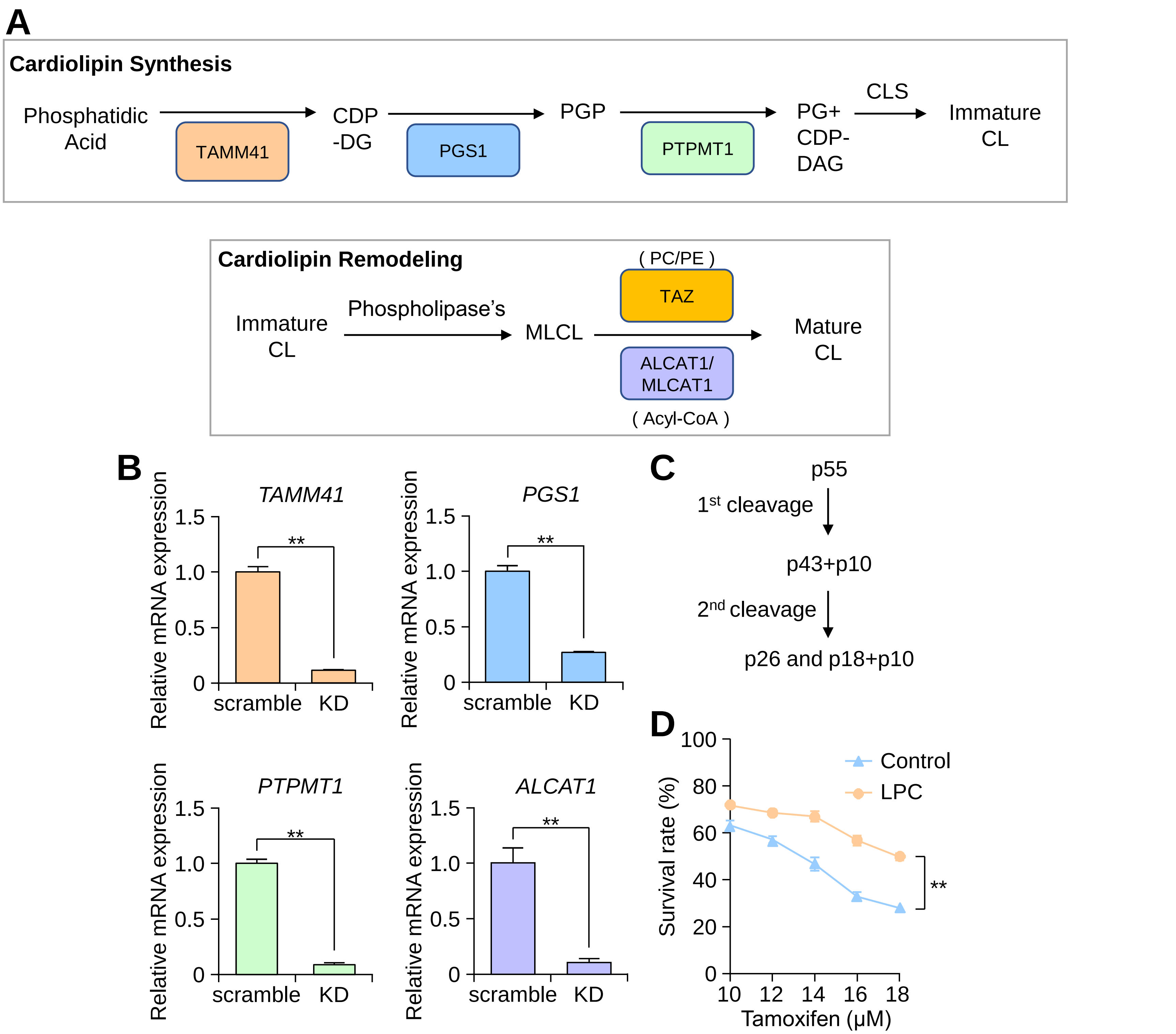

### Supplemental Figure S6

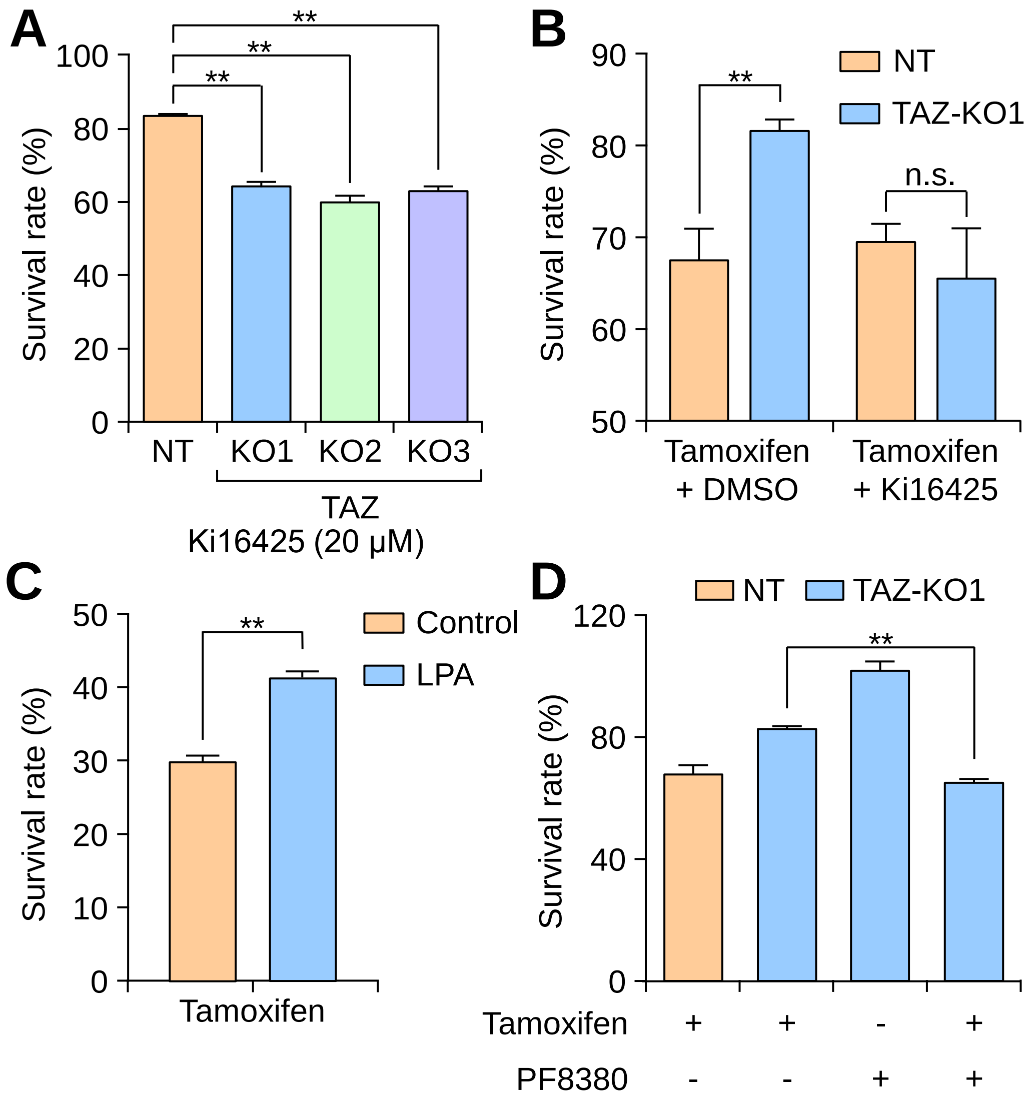
