## Supplemental Table 1 for "Tafazzin Mediates Tamoxifen Resistance by Regulating Cellular Phospholipid Composition in ER-Positive Breast Cancer"

**Table S1. sgRNA sequences used for validation of tamoxifen-resistance candidate genes from the initial screen, related to Methods.**

| Target Gene | sgRNA sequence (5‘-3’) |
| --- | --- |
| Non-Targeting | AAAAAGCTTCCGCCTGATGG |
| TAZ-sg1 | CCTGACCGTGCACAACAGGG |
| TAZ-sg2 | GAATTCCTGCGTTTCAAGTG |
| TAZ-sg3 | GAGATGGCGTCTACCAGAAG |
| RPS16-sg1 | ACAGAAGACAGCGACAGCTG |
| RPS16-sg2 | ATCCGTGTCCGTGTAAAGGG |
| PELO-sg1 | ACTGCTCCCTCACAAATCCT |
| PELO-sg2 | TGACTAAGCAGATATGGGCG |
| NDUFB4-sg1 | CATATCTCCGGAAACCCGGC |
| NDUFB4-sg2 | CAGGTACTCTCGTTTCAGCT |
| IMP3-sg1 | GCGCAGCTTGAGGAGCACGG |
| IMP3-sg2 | AGCTCCAGCGAACCGCGCGT |
| ROMO1-sg1 | ACGGTCGAAGCAGCTTGGCT |
| ROMO1-sg2 | CTCCTCAGGATCGGAATGCG |
| PSMD13-sg1 | ATGAAGAATGATTTCCACGA |
| PSMD13-sg2 | GAACCGATGTCACACCAGGA |
| USPL1-sg1 | CCTAATGTGCATCTAAGCTG |
| USPL1-sg2 | GTCCTGTGAAAGAGTACGAG |
| CCDC81-sg1 | AGCTTGCCAGGATCATAACA |
| CCDC81-sg2 | TTCTCCACCATGATAAACAC |
