## Supplemental Table 2 for "Tafazzin Mediates Tamoxifen Resistance by Regulating Cellular Phospholipid Composition in ER-Positive Breast Cancer"

**Table S2. sgRNA sequences for correlation analysis of cell proliferation index and tamoxifen resistance index, related to Figure 3.**

| Gene symbol | sgRNA sequence (5‘-3’) | Proliferation  Index | Resistance  Index |
| --- | --- | --- | --- |
| Low-proliferation | | | |
| MTOR (sg1) | GGTGATGGCCTGGACAACCA | 0.15 | 1.57 |
| PSMB2 | TTTATAAGATGCGAAATGGT | 0.23 | 1.68 |
| WASH1 | GGTGGTGTTGAAGAGCAGCA | 0.26 | 1.38 |
| EIF3CL | GCAGAGCATGGTGGTAGATG | 0.30 | 2.35 |
| MTOR (sg2) | TCAGGAAATGATCCGCACAG | 0.31 | 1.43 |
| PPP1R7 | GCTGGGATCTAACCGCATCC | 0.32 | 0.75 |
| RPS12 | TCCACCAACTTGACATACAT | 0.34 | 1.13 |
| NOTCH2NL (sg1) | GGGATGAGACAGGCAGGCAT | 0.35 | 1.30 |
| EXOSC8 | GGAAATACTACAGTAATCTG | 0.35 | 1.63 |
| SOD1 | GGAAAGTAATGGACCAGTGA | 0.38 | 1.49 |
| PSMD13 | ATGAAGAATGATTTCCACGA | 0.41 | 1.79 |
| USP7 | GGCAACCTTTCAGTTCACTG | 0.47 | 1.69 |
| NF1 | GAGGAAGCAGATATCCGGTG | 0.47 | 1.56 |
| SARS2 | TACCGGGCAGAGACAAACAC | 0.48 | 2.10 |
| Medium-proliferation | | | |
| NOTCH2NL (sg2) | TCTCGACCTTGCCTGAATGG | 0.51 | 0.98 |
| EXOC4 | TTCTCATAGATGAACTACAC | 0.51 | 1.60 |
| PIK3C3 | AGCCTGCAAAAACTCAACAC | 0.57 | 1.19 |
| USPL1 | CCTAATGTGCATCTAAGCTG | 0.57 | 1.24 |
| MOCS3 | AATCTCATCTCGGGACAGAG | 0.63 | 1.51 |
| GATA1 | GCGGGTGGGACACACAGTTG | 0.68 | 0.98 |
| AURKA | CCATATAGAAAATAATCCTG | 0.72 | 1.09 |
| CIB3 | AAGGTGCCCTACGAGCTCAT | 0.74 | 0.93 |
| CGB1 | GCATTGATGGGGCGGCACCG | 0.76 | 0.97 |
| NOL7 | GGCGAAAGTCAGCTCCTCCG | 0.80 | 1.40 |
| TAZ | CCTGACCGTGCACAACAGGG | 0.88 | 2.13 |
| LATS1 | CCTTCTGCTTTACAAACAGG | 0.89 | 1.18 |
| RAB9B | CTTAGGACACCCTTCTACAG | 0.92 | 1.12 |
| CCDC81 | AGCTTGCCAGGATCATAACA | 0.98 | 0.90 |
| High-proliferation | | | |
| CNNM3 | ATGGTTGTAGAAACGAGTGA | 1.24 | 1.33 |
| GOLPH3 | TGTATATCATCTGGATTACG | 1.37 | 1.16 |
| MRAS | GGAGCAATACATGCGCACGG | 1.50 | 1.24 |
