## Supplemental Table 3 for "Tafazzin Mediates Tamoxifen Resistance by Regulating Cellular Phospholipid Composition in ER-Positive Breast Cancer"

**Table S3. Primers used for qPCR and PCR gene cloning, related to Methods.**

| Gene | Forward (5‘-3’) | Reverse (5‘-3’) |
| --- | --- | --- |
| qPCR primers | | |
| *GAPDH* | GGAGCGAGATCCCTCCAAAAT | GGCTGTTGTCATACTTCTCATGG |
| *TAZ* | CACCGTGTCCAATCACCAGTC | TCCAACGCATCAACTTCAGGT |
| *CCND1* | GCTGCGAAGTGGAAACCATC | CCTCCTTCTGCACACATTTGAA |
| *ESR1* | GGGAAGTATGGCTATGGAATCTG | TGGCTGGACACATATAGTCGTT |
| *GREB* | ATGGGAAATTCTTACGCTGGAC | CACTCGGCTACCACCTTCT |
| *MYC* | GGCTCCTGGCAAAAGGTCA | CTGCGTAGTTGTGCTGATGT |
| *PGR* | TTATGGTGTCCTTACCTGTGGG | GCGGATTTTATCAACGATGCAG |
| *TFF1* | CCCCGTGAAAGACAGAATTGT | GGTGTCGTCGAAACAGCAG |
| *TFF2* | GCTGTTTCGACTCCAGTGTCA | CCACAGTTTCTTCGGTCTGAG |
| *TFF3* | CCAAGCAAACAATCCAGAGCA | GCTCAGGACTCGCTTCATGG |
| *TAMM41* | CAGTAGATGACCCTGTCGCAT | GGATGGACGTGATAATCTTGGG |
| *PGS1* | TCGTGATGGCATCCCTCTAC | CCCCGCGTGAAGTCTAAGA |
| *PTPMT1* | CAGAGGAGGCTGTAAGAGCCA | TGTGGATGTATGACCGGATCT |
| *ALCAT1* | CACCCTACCTGTGGCATTATTG | CCATTCTTGTCCGATGGTTCAT |
| *ATX* | ACTTTTGCCGTTGGAGTCAAT | GGAGTCTGATAGCACTGTAGGA |
| PCR primers | | |
| TAZ | ATGCCTCTGCACGTG | CTATCTCCCAGGCTGGAG |
| ESR1 | ATGACCATGACCCTCCACACC | TCAGACCGTGGCAGGGAAACCC |
