## Supplemental Table 4 for "Tafazzin Mediates Tamoxifen Resistance by Regulating Cellular Phospholipid Composition in ER-Positive Breast Cancer"

**Table S4. shRNA sequences used in the study, related to Methods.**

| Target Gene | shRNA sequence (5‘-3’) |
| --- | --- |
| Scramble | CCTAAGGTTAAGTCGCCCTCG |
| TAZ-sh1 | CGGACTTCATTCAAGAGGAAT |
| TAZ-sh2 | CTGTACGAGCTCATCGAGAAG |
| TAMM41 | GACTTTGTGTTCACAGTAGAT |
| PGS1 | GCCTGGAAAGTACTCTAGAAA |
| PTPMT1 | GATCCGGTCATACATCCACAT |
| ALCAT1 | CAGTCTTGTTAAGTGGTATTT |
