## Supplemental Table 5 for "Tafazzin Mediates Tamoxifen Resistance by Regulating Cellular Phospholipid Composition in ER-Positive Breast Cancer"

**Table S5. TIDE analysis of candidate genes knockout cells.**

| Gene | Wt | Ko | Efficiency |
| --- | --- | --- | --- |
| PELO-sg1 | 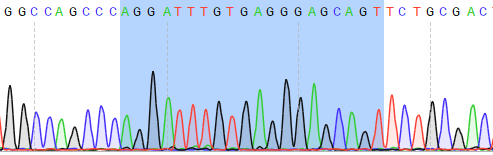 | 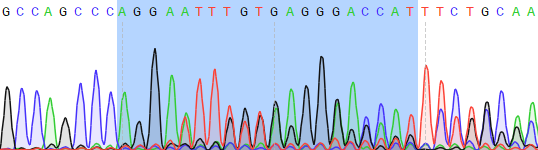 | 63.6 |
| PELO-sg2 | 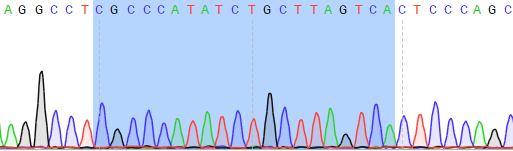 | 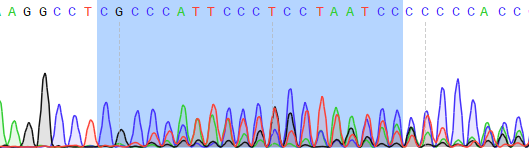 | 35.4 |
| ROMO1-sg1 | 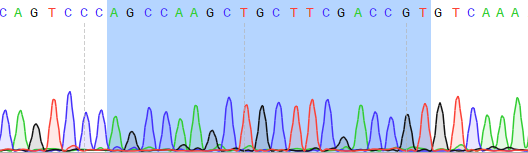 | 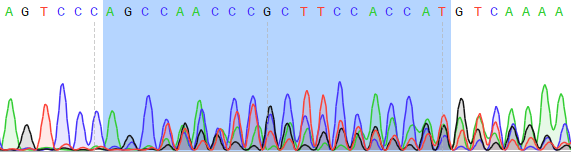 | 35.2 |
| ROMO1-sg2 | 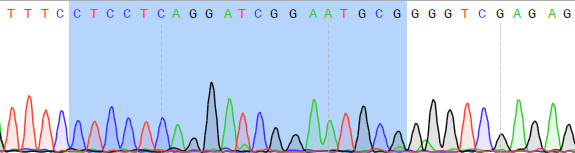 | 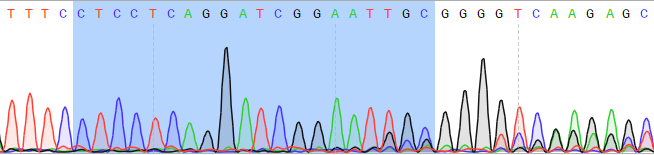 | 9.9 |
| TAZ-sg1 | 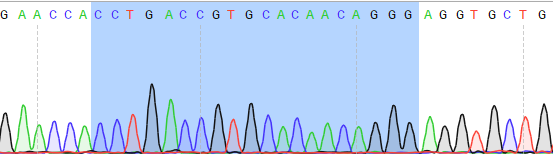 | 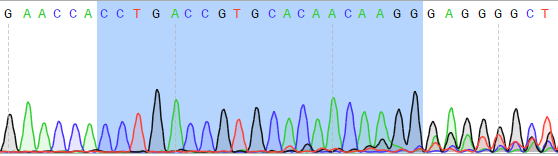 | 76.9 |
| TAZ-sg2 | 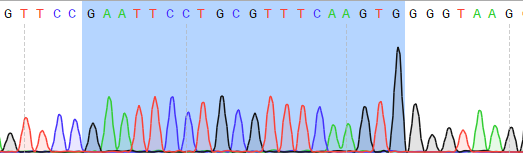 | 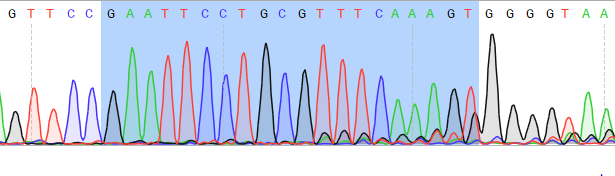 | 88.8 |
| USPL1-sg1 | 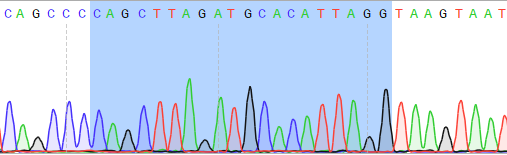 | 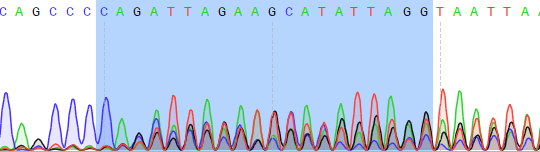 | 49 |
| USPL1-sg2 | 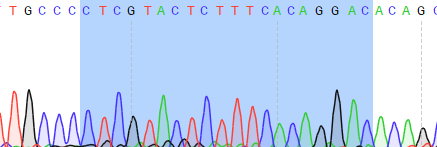 | 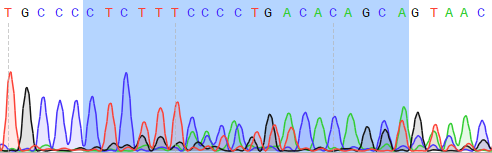 | 85.9 |
| CCDC81-sg1 | 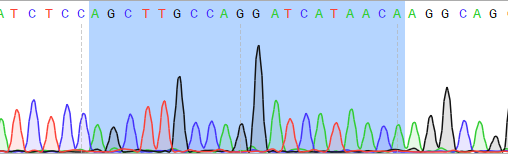 | 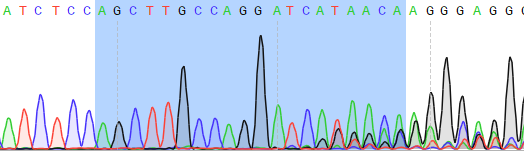 | 18.5 |
| CCDC81-sg2 | 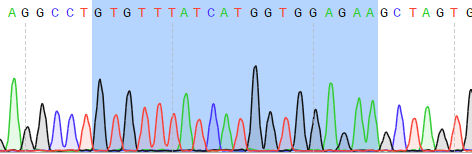 | 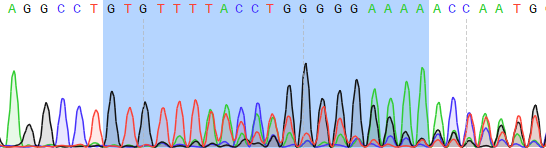 | 57.2 |
| IMP3-sg1 | 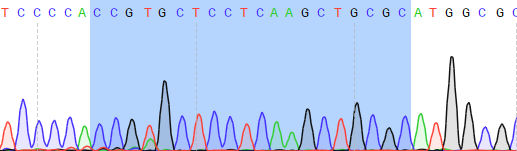 | 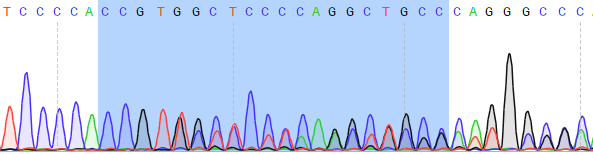 | 61.7 |
| IMP3-sg2 | 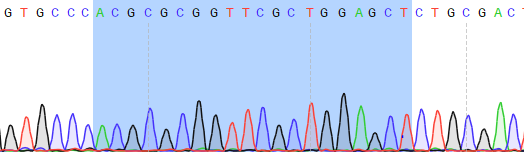 | 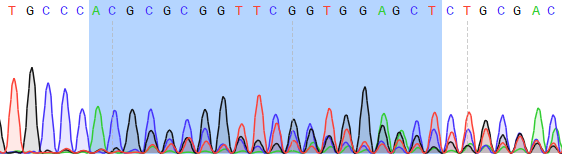 | 48.4 |
| PSMD13-sg1 | 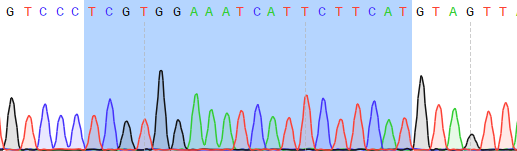 | 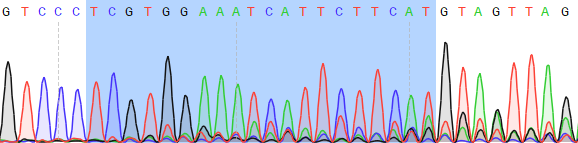 | 34.7 |
| PSMD13-sg2 | 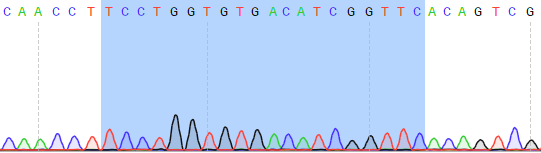 | 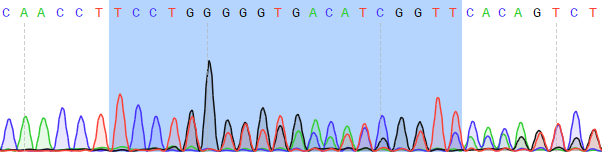 | 33.1 |
| NDUFB4 | 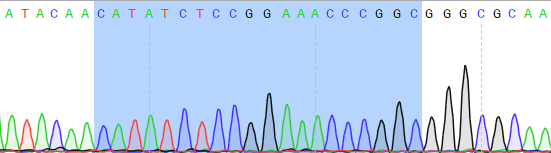 | 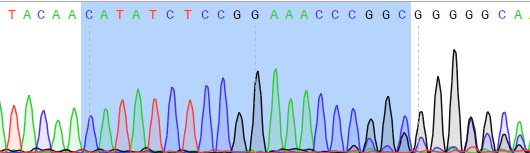 | 52.6 |
